# Trait-based life-history strategies explain succession scenario for complex bacterial communities under varying disturbance

**DOI:** 10.1101/546416

**Authors:** Ezequiel Santillan, Hari Seshan, Florentin Constancias, Stefan Wuertz

**Author notes:** Correspondence to: Stefan Wuertz,. Present address: Brown and Caldwell, 9665 Chesapeake Drive, Suite 201, San Diego CA 92123, U.S.A.

## Abstract

Trait-based approaches are increasingly gaining importance in community ecology, as a way of finding general rules for the mechanisms driving changes in community structure and function under the influence of perturbations. Frameworks for life-history strategies have been successfully applied to describe changes in plant and animal communities upon disturbance. To evaluate their applicability to complex bacterial communities, we operated replicated wastewater treatment bioreactors for 35 days and subjected them to eight different disturbance frequencies of a toxic pollutant (3-chloroaniline), starting with a mixed inoculum from a full-scale treatment plant. Relevant ecosystem functions were tracked and microbial communities assessed through metagenomics and 16S rRNA gene sequencing. Combining a series of ordination, statistical and network analysis methods, we associated different life-history strategies with microbial communities across the disturbance range. These strategies were evaluated using tradeoffs in community function and genotypic potential, and changes in bacterial genus composition. We further compared our findings with other ecological studies and adopted a semi-quantitative CSR (*competitors, ruderals, stress-tolerants*) classification. The framework reduces complex datasets of microbial traits, functions, and taxa into ecologically meaningful components to help understand the system response to disturbance, and hence represents a promising tool for managing microbial communities.

**Originality-Significance Statement:** This study establishes, for the first time, CSR life-history strategies in the context of bacterial communities. This framework is explained using community aggregated traits in an environment other than soil, also a first, using a combination of ordination methods, network analysis, and genotypic information from shotgun metagenomics and 16S rRNA gene amplicon sequencing.

## Introduction

Biogeochemical cycles are primarily driven by microbial activity (Widder *et al*. 2016), and the ever increasing anthropogenic impact on the biosphere calls for a better understanding of the mechanisms that structure microbial communities in response to human-induced disturbances (Falkowski *et al*. 2008). Disturbance in general is deemed a major factor influencing variations in species diversity (Mackey and Currie 2001) and structuring of ecosystems (Shade *et al*. 2012a, Shade *et al*. 2012b). Additionally, unravelling what drives patterns of community succession and structure remains a central goal in ecology (Powell *et al*. 2015, Zhou *et al*. 2014), especially since community diversity is thought to regulate community function (Mouillot *et al*. 2013). The challenge of finding general rules in community ecology (Lawton 1999, Simberloff 2004) has fuelled a growing interest in the role of functional traits in community ecology (McGill *et al*. 2006, Westoby and Wright 2006). Such traits are defined as morphologic, physiologic, genomic or phenotypic attributes that affect the fitness (growth, reproduction and survival) of an organism (Violle *et al*. 2007). Organisms face tradeoffs to allocate resources into certain traits to maximize their fitness, which depends on abiotic and biotic interactions within the environment they inhabit. Trait-based approaches in community ecology have been applied to eukaryotic microbial communities like phytoplankton (Edwards *et al*. 2011, Litchman and Klausmeier 2008, Litchman *et al*. 2007) and fungi (Chagnon *et al*. 2013, Crowther *et al*. 2014, Treseder and Lennonb 2015). More recently, trait-based approaches have been proposed for the study of bacterial community dynamics to link biodiversity and ecosystem functioning, with an emphasis on tradeoffs (Krause *et al*. 2014) and phylogenetic conservation of traits (Martiny *et al*. 2013, Martiny *et al*. 2015). Given the challenges of defining species for prokaryotes (Rossello-Mora and Amann 2015), there is a trend to shift the focus from ‘who are they’ to ‘how will they respond’ (Boon *et al*. 2014). In this regard, ecological theory can help to elucidate the underlying mechanisms structuring communities in studies of complex microbial dynamics (Prosser *et al*. 2007).

The theoretical CSR framework for plant communities developed by Grime (1977) constitutes a classic trait-approach and proposes three types of life-history strategies: competitors (C) who maximize resource acquisition and control in consistently productive niches, stress-tolerants (S) who can maintain metabolic performance in unproductive niches, and ruderals (R) who have good growth rates but inefficient resource uptake in niches where events are frequently detrimental to the individual. Such strategies depend on varying intensities of disturbance (biomass destruction), stress (biomass restriction), and competition for resources (Fig. S1). The application of the CSR framework has been expanded from plants to other organisms (Grime and Pierce 2012) and it has proven to be a useful tool for conservation and ecosystem management (Grime 2013), despite some criticism (Wilson and Lee 2000). The CSR framework was recently suggested for microbial communities in a series of reviews that gathered data from different studies and identified traits and tradeoffs to classify different methane-oxidizing bacteria (Ho *et al*. 2013) and arbuscular mycorrhizal fungi (Chagnon *et al*. 2013). Further reviews emphasized the potential of employing the CSR as a trait approach to anticipate changes in microbial community structure during succession, proposing some examples of traits that could be generally related to each life strategy (Crowther *et al*. 2014, Ho *et al*. 2017, Krause *et al*. 2014). However, meta-analyses of microbial communities to infer CSR strategies can be challenging due to the different factors related to experimental design and techniques employed across different studies, which could heavily affect the observations in comparison. Recent studies on soil-bacterial communities in cadmium-contaminated rhizospheres (Wood *et al*. 2018) and tillage-disturbed fields (Schmidt *et al*. 2018) also suggested the applicability of the CSR approach to soil-microbial communities. To our knowledge, the CSR framework has not yet been employed within a single study to changes in microbial communities upon disturbance in any microbiome other than soil.

Analysing tradeoffs in traits and functions on the level of a whole community by identifying community aggregated traits (CATs) (Shipley *et al*. 2006) represents an additional opportunity to tackle the complexity of microbial community dynamics. Community-level traits arise from an array of diverse organisms interacting in a direct and indirect way under environmental gradients, resulting in an overall ecosystem function that can greatly differ from the ecological attributes of individual taxa (Fierer *et al*. 2014). Beyond measuring ecosystem function, metagenomics data can be used to infer CATs assuming that the metagenomes represent a random sampling of all microbial genomes, from which we can gather genotypic traits of interest (Barberan *et al*. 2012). Based on CATs, the CSR life-history strategies framework could be applied to whole communities, to identify life-history strategies beyond specific taxa. This has not been assessed yet in any study of microbial communities.

Here we investigate the effect of disturbance in bacterial community structure, genotypes and function within a framework of three-way CSR life-history strategies. We conducted a 35-day laboratory study using sequencing batch bioreactors inoculated with activated sludge from an urban wastewater treatment plant. The experiment involved different frequency levels of augmentation with toxic 3-chloroaniline (3-CA) as disturbance, with triplicate reactors receiving 3-CA either never (L0, undisturbed), every seven, six, five, four, three, and two days (L1-6, intermediately-disturbed), or every day (L7, press-disturbed). Sludge reactors are model systems for microbial ecology (Daims *et al*. 2006), harbouring complex microbial communities with defined and measurable ecosystem functions (Seviour and Nielsen 2010). Chloroanilines are xenobiotic, carcinogenic and toxic substances, known to hinder both nitrogen and carbon removal in sludge bioreactors (Falk and Wuertz 2010). A microcosm design allowed us to test a wide range of disturbances with good replication, while minimizing confounding factors across replicates (Drake and Kramer 2012). Our approach combined analyses of variations in ecosystem function, bacterial composition and abundances, and genotypic traits across a wide gradient of disturbances. We hypothesized that the extreme sides of our proposed disturbance range would favour two distinct life-history strategies (competitors, C, and stress-tolerants, S), while intermediate levels of disturbance will harbour a gradient of ruderal (R)-type strategies. We further hypothesized that such differentiation in life-history strategies would be attributable at the whole bacterial community level.

## Results

### Patterns of community structure and functional tradeoffs across disturbance

Bacterial communities differentiated in terms of β-diversity across disturbance levels and with time, as revealed by community analysis through 16S rRNA amplicon sequencing. There was a temporal separation of communities at the undisturbed level 0, the intermediately disturbed levels 1 to 6, and the press-disturbed level 7 as shown by unconstrained ordination (Fig. 1a). Furthermore, differences in community structure assessed through PCR-independent shotgun metagenomics were marked across different disturbance levels as shown by constrained ordination on d35. The primary axis (Fig. 1b) differentiates the intermediately disturbed levels (L1-6) from the press-disturbed one (L7), while the secondary axis highlights the separation of the undisturbed level (L0). Multivariate tests yielded significant results for disturbance levels (PERMANOVA-P = 0.0003), but with a significant effect of heteroscedasticity (PERMDISP-P = 0.014). However, GLMMs tests after fitting data to a negative-binomial distribution yielded significant results (P = 0.015), confirming that the observed differences among groups were due to disturbance levels and not only to heteroscedasticity.

**Fig. 1.**
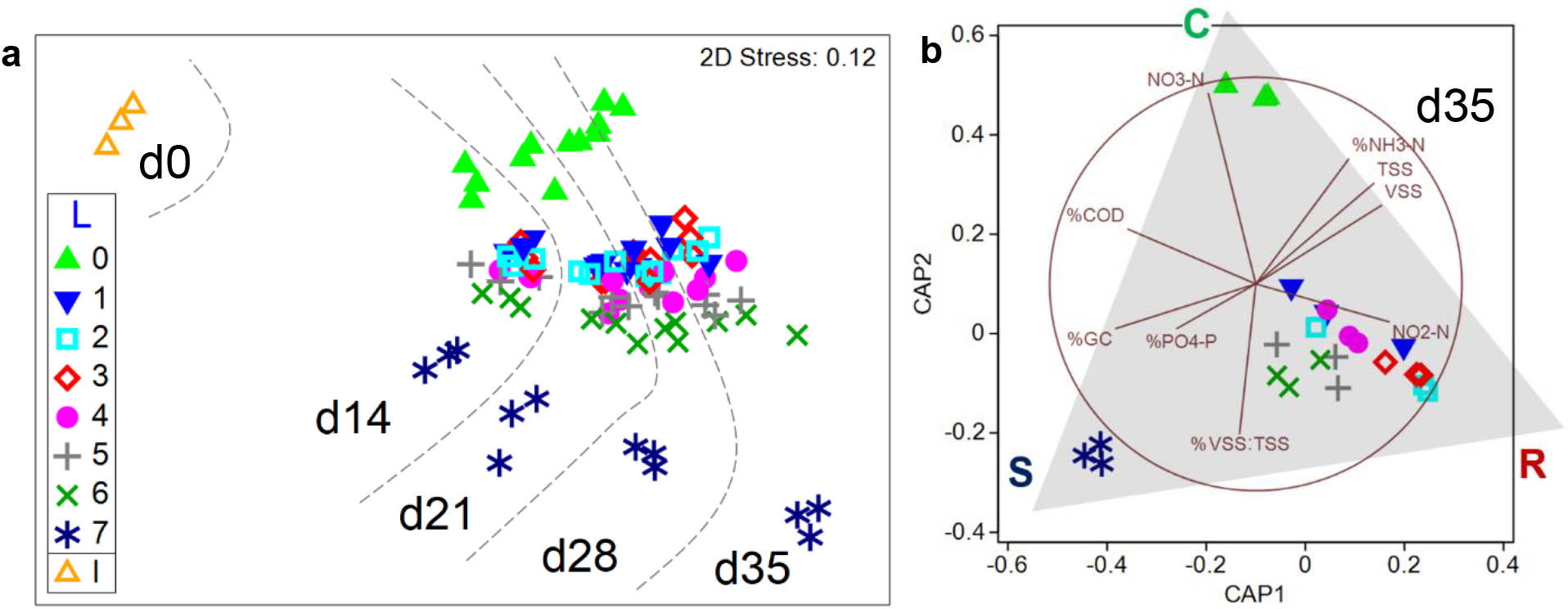
Bacterial community structure variations with time and disturbance. (**a**) Community succession in terms of β-diversity patterns, assessed through non-metric multidimensional scaling (NMDS) of 16S rRNA gene amplicon sequence variant level data. Dashed lines indicate different days. (**b**) Canonical analysis of principal coordinates (CAP) on shotgun genus-level metagenomics data at d35. Pearson’s correlation vectors (r > 0.40) represent community-level functions, highlighting tradeoffs in soluble organic carbon (%COD), ammonia (%NH_3_-N), and phosphorus removal (%PO_4_-P); nitrite (NO_2_-N) and nitrate production (NO_3_-N); GC content in metagenomic DNA (%GC); total (TSS) and volatile solids (VSS); and biomass fraction (%V/T). Shaded triangle overlaid to highlight CSR life-history strategies: C, competitors (L0); R, ruderals (L1-6); S, stress-tolerants (L7). Legends: L, disturbance levels 0-7; I, full-scale plant sludge inoculum.

Variations in β—diversity corresponded with tradeoffs in community-level function, indicated by Pearson’s correlation vectors (Fig. 1b). Ecosystem function parameters measured across reactors at different levels of disturbance were significantly different (Table S1). Undisturbed reactors (L0) displayed maximum soluble chemical oxygen demand (COD) and NH_3_-N removal, and maximum NO_3_-N production without any detectable NO_2_-N residual. At the other extreme, press-disturbed reactors (L7) had a high COD removal, but the worst NH_3_-N removal with no detectable NO_X_-N concentrations. Intermediately disturbed levels (L1-6) displayed variable NO_2_-N effluent concentrations across replicates, which decreased with increasing disturbance. The maximum values of NO_2_-N coincided with minimum COD removal, suggesting a tradeoff between such functions under disturbance. NO_3_-N production for L1-6 was minimal, but with a similar trend than for NO_2_-N. Total (TSS) and biomass (VSS) values were lowest for the press-disturbed level (L7) and highest for the lowest disturbance level (L1). Differences in PO_4_-P removal were not significant, despite being 20-30% higher on average for L6 and L7. All disturbed reactors (L1-7) showed complete 3-CA degradation, a metabolic capacity that was initially absent and later acquired by the community during the experiment (see Santillan *et al*. (2019) for detailed information of temporal ecosystem function changes).

### Genera abundances are differentially distributed across the disturbance range

Relative abundance comparisons revealed diverse bacterial taxa prevailing at different levels of disturbance, as shown for the 25 most abundant genera after 35 days assessed through metagenomics (Fig. 2). Genera like *Nitrospira, Paracoccus*, and *Dehalobacter* were enriched in the undisturbed treatment (L0), while *Gemmatimonas* and *Mesorhizobium* were favoured at the press-disturbed level (L7). Other organisms like *Nitrosomonas, Labilithrix* and *Nakamurella* benefited at intermediate levels of disturbance (L1-6). *Tetrasphaera* and *Pyrinomonas* decreased in abundance with disturbance, while the opposite trend was found for taxa like *Thauera* and *Ca.* Contendobacter. Extreme conditions of no disturbance (L0) and press disturbance (L7) favoured *Microlunatus* and *Bosea*. Genera clusters were evident across the disturbance range in a heat map analysis of the top 100 genera (Fig. S2).

**Fig. 2.**
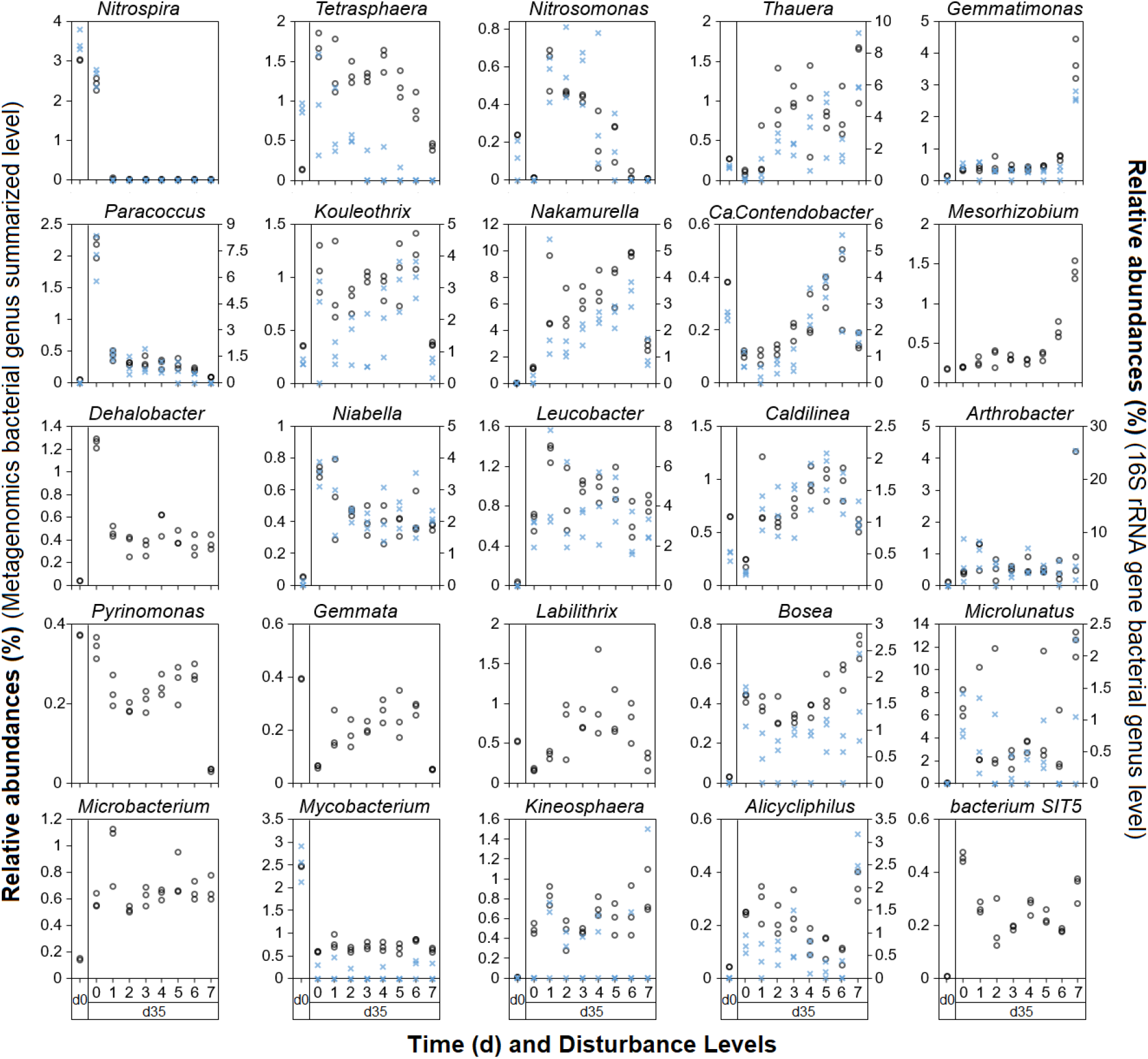
Relative abundance comparisons to discern taxa prevailing at different disturbance levels across reactors (n = 24). Top 25 genera assessed through metagenomics (open circles, left y-axis) at d35. Relative abundances at d0 included for comparison with initial conditions. Relative abundances from 16S rRNA gene amplicon sequencing are included for detected genera (blue crosses, right y-axis when needed).

Most of the top 25 metagenomics genera were also detected through 16S rRNA gene amplicon sequencing, showing similar compositional trends across disturbance levels (Fig. 2). For some genera, like *Nitrospira* and *Gemmatimonas*, the relative abundances nearly coincided for both sequencing methods. The ranking of the top 25 genera detected by 16S rRNA amplicon sequencing (Fig. S3) differed from that detected by metagenomics, but similar abundance patterns across disturbance levels were observed. Taxa like *OLB17, Paraccocus*, and *env.OPS_17* prevailed at the undisturbed level (L0); while others like *OLB1, Nakamurella* and *Plasticicumulans* did so at intermediately-disturbed levels (L1-6). *Thauera* and *SBR1031* increased their abundances with disturbance, while *Actinomycetaceae* and *Saccharimonadales* were favoured at undisturbed (L0) and press-disturbed (L7) levels. Changes in relative abundances of bacterial genera across time and disturbance levels were marked for all reactors (Fig. S4). Still, several relevant genera (i.e., *Nitrospira, Nitrosomonas, Kouleothrix, Ca.* Contendobacter) had comparable relative abundances to that of the full-scale plant inoculum (d0) for at least one of the disturbance levels assessed on d35 (Fig. 2, Figs. S2-4).

### Different genotypic traits favoured with varying disturbance

Functional analysis of the metagenomics dataset revealed groups of genotypic traits associated with different levels of disturbance. In order to capture broad changes favoured by different life-history strategies we focused on trait complexes, which are a product of the expression of multiple true traits (Crowther *et al*. 2014), grouping different sets of genes into categories. A double-clustered heat map, including 57 trait complexes mapped from the InterPro2Go (IP2G) database (Finn *et al*. 2017), showed that replicate reactors at the undisturbed (L0) and press-disturbed level (L7) formed two separated clusters (Fig. 3). The differential prevalence of community-aggregated genotypic traits (Fierer *et al*. 2014) provided additional support for different life-history strategies across the disturbance range. Following the CSR framework, we classified these traits as C, CR, R, SR, S, or CS according to z-score values ≥ +1.0 on the heat map (see Fig. S5 for relative abundances). Only differences in the gene traits of cell communication, hydrolase activity, and enzyme regulator activity were found not to be significant after univariate testing (Table S2). Similar clustering trends emerged for trait complexes obtained from additional functional databases like COG, KEGG and SEED (Fig. S6).

**Fig. 3.**
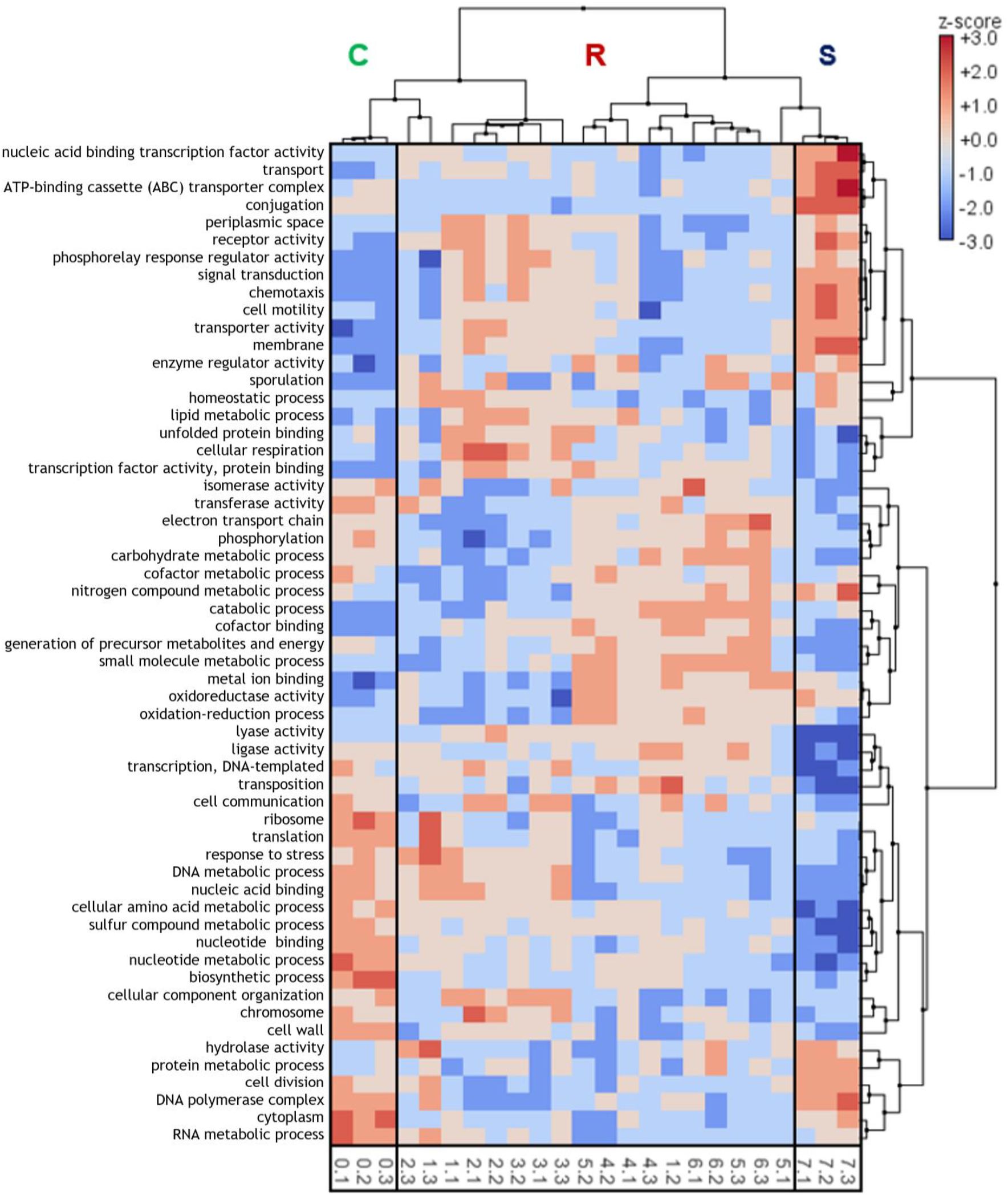
Heat map of genotypic potential traits after mapping metagenomes with the IP2G database (Gene Ontologies) at d35. Functional capacity is classified as trait complexes of biological processes, cellular component, and molecular function. Only traits with more than 10,000 reads assigned across all 24 reactors were considered. Clustering was applied to differentiate groups of trait complexes and disturbance levels. Rectangles highlight taxa groups prevailing at different CSR life-history strategies at the community-level: C, competitors (L0); R, ruderals (L1-6); S, stress-tolerants (L7).

The relationships among observed community structure patterns and different community-level genotypes across disturbance levels was assessed using distance-based redundancy analysis (dbRDA) using the IP2G trait complexes as predictor variables (Fig. 4). Up to 71.4% of the variance could be fitted in the first two axes of the dbRDA, which is considered a good model fit (Clarke *et al*. 2014). Significant Pearson’s correlations highlighted some traits favoured by bacterial communities at different levels of disturbance (Fig. 4). Furthermore, correlation-based network analysis showed comparable clusters based on node modularity for bacterial taxa and genes at different levels of resolution. There were similar clusters for the top 200 metagenomics genera and top 57 trait complexes from the IP2G database (Fig. 5), as well as for the top 200 16S rRNA gene amplicon sequence variants and top 200 individual genes from the IP2G database (Fig. S7).

**Fig. 4.**
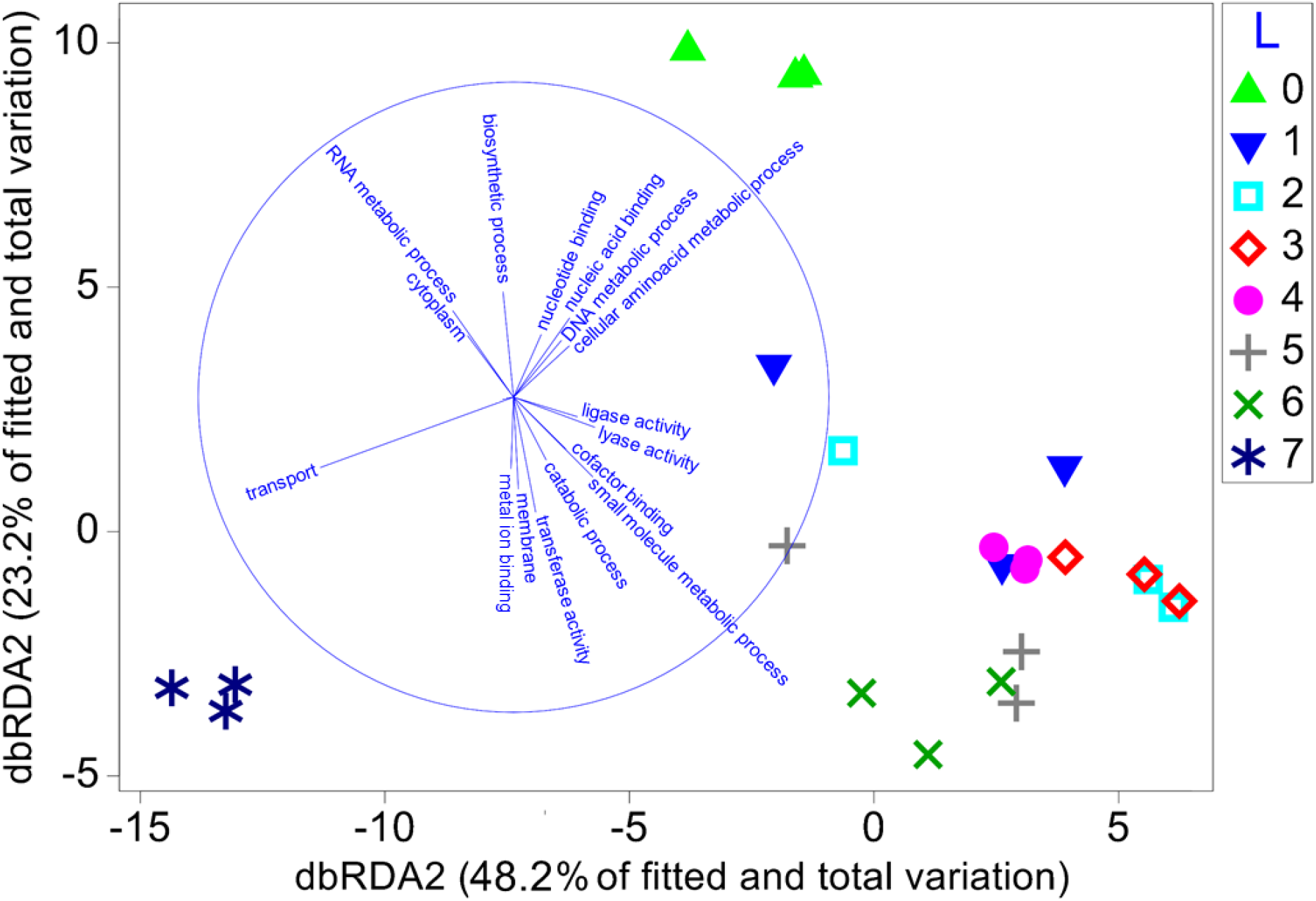
Bacterial genotypic traits relationship with community structure across disturbance. Constrained ordination of distance-based redundancy analysis (dbRDA) using trait complexes of the IP2G database (gene ontologies) as predictor variables of metagenomics genus-level community data at d35. Pearson’s correlation vectors (r > 0.20) represent some of the trait complexes favoured by different bacterial communities.

**Fig. 5.**
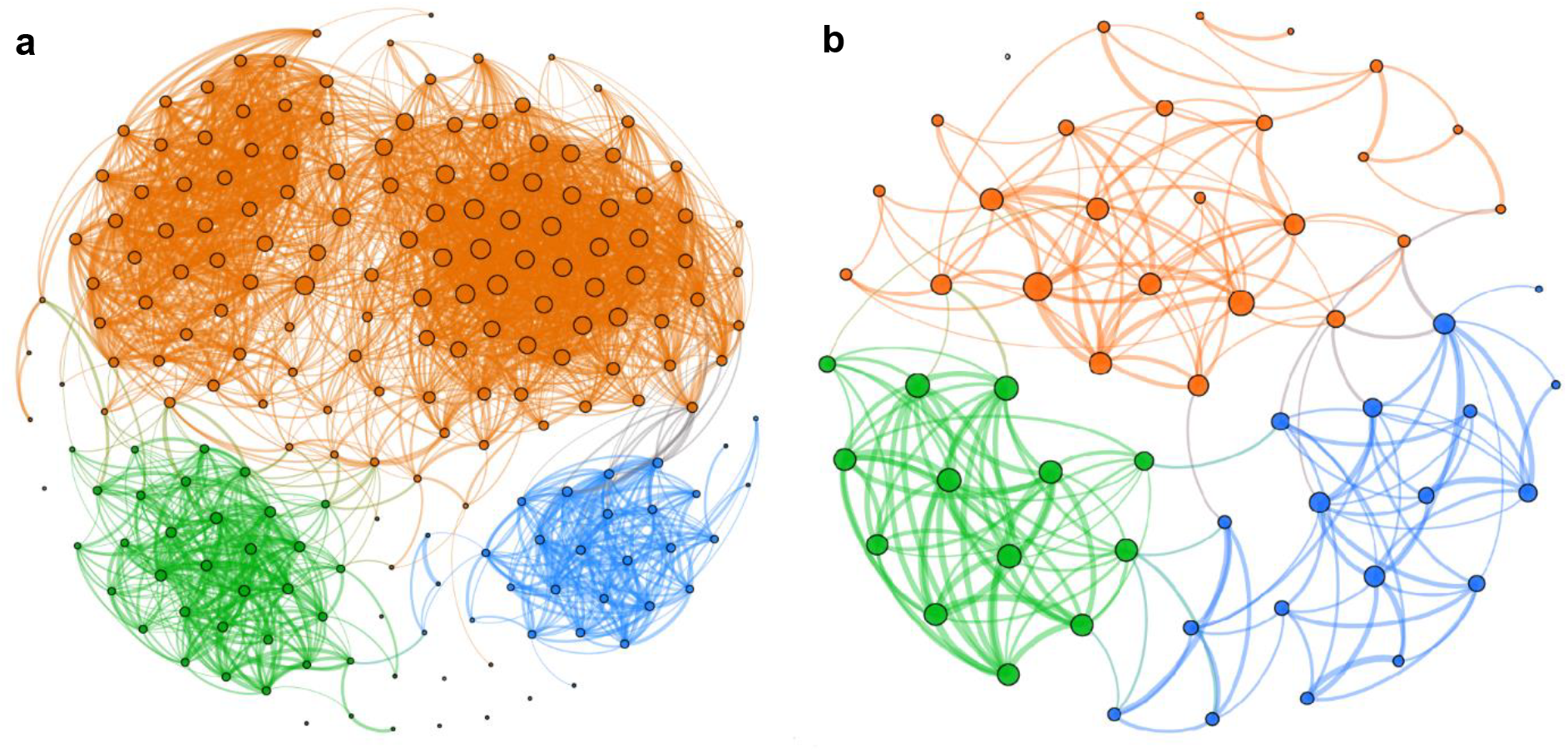
Correlation networks display separated clusters based on node modularity for bacterial taxa and genes. (**a**) Top 200 metagenomics genera. (**b**) Top 57 genotypic categories (trait complexes) from IP2G database. Clusters are coloured by modularity class, with green nodes prevailing in undisturbed (L0) reactors, orange nodes in intermediately disturbed reactors (L1-6), and blue nodes in press-disturbed reactors (L7). Only significantly strong Pearson’s correlations (r ≥ 0.50) were employed. For each panel, edge thickness represents correlation strength and node size represents degree.

## Discussion

### Disturbance, succession, and community-level life-history strategies

We employed sludge for wastewater treatment as a model system (Daims *et al*. 2006), under a successional study in which the inoculum community was taken from a full scale plant and then used to seed our microcosm bioreactors. The experiment purposely lacked an acclimation phase, hence the following alterations were expected to promote succession within microbial communities: feeding scheme (continuous to batch), volume (full scale to microcosm), cell residence time (low to high), feed type (natural to synthetic wastewater), immigration (open to closed system). Furthermore, the effect of disturbance within succession was assessed by varying frequencies of toxic 3-CA included in the synthetic feed for the reactors. Indeed, bacterial community succession was observed through 16S rRNA sequencing analysis, in terms of temporal patterns of β-diversity (Fig. 1a) and genus-level relative abundances (Fig. S4). Metagenomics data also displayed changes in bacterial community structure between the inoculum (d0) and all reactors at d35 (Fig. 2, Fig. S2). We expected to observe changes in community structure after 35 days, considering that activated sludge contains bacteria with generation times varying from 20 minutes to days, with most relevant bacteria doubling in less than 24 hours (Tchobanoglous *et al*. 2003). Hence, this study constituted an appropriate scenario for the CSR framework, as one of its main applications lies in predicting community shifts during succession after disturbance (Caccianiga *et al*. 2006, Grime and Pierce 2012).

There were clear differences among sludge reactors that had been exposed to varying disturbance frequencies, in terms of community structure, function, and genotypic traits on d35. We therefore propose that microbial communities adopted different CSR life-history strategies under the disturbance regime imposed in our study, with undisturbed L0 as *competitors*, the press-disturbed L7 as *stress-tolerants*, and the remaining intermediately disturbed levels as *ruderals*. Community-level functional tradeoffs across life-history strategies were marked: organic carbon removal was higher for competitors and stress-tolerants; complete nitrification was only achieved by competitors; incomplete (inefficient) nitrification prevailed among ruderals; and biomass productivity was higher for ruderals and competitors than for stress-tolerants (Table S1).

There were different community-level (Fierer *et al*. 2014) trait complexes (Crowther *et al*. 2014) prevailing at different levels of disturbance, which also showed a three-way CSR clustering by means of network analysis (Fig. 5, Fig. S7) and heat maps (Fig. 3, Figs. S2 and S6). It was also evident by means of two different sequencing techniques (Knight *et al*. 2018), that different bacterial genera were favoured at different level of disturbance (Fig. 2, Figs. S2-4). Furthermore, ordination methods showed CSR-like β-diversity distinctions among communities across disturbance levels. Different ordination methods for species and trait abundances datasets have been widely employed in ecology to identify and validate three-way life-history strategies. For example, a combination of DCA, NMDS, and PCA was employed to describe patterns of grass vegetation succession in a glacier foreland (Caccianiga *et al*. 2006); a meta-study on reef corals used PCO (Darling *et al*. 2012); and PCA was used on fish survey datasets (Pecuchet *et al*. 2017). We found that bacterial communities grouped in three major β-diversity clusters using NMDS and CAP ordinations on datasets arising from different sequencing techniques (Fig. 1). Overlaid correlation vectors of process performance highlighted tradeoffs in ecosystem function on d35 across those clusters (Fig. 1b). Likewise, a distance-based redundancy analysis showed a relationship for community β-diversity patterns and genotypic traits (Fig. 4).

### Combining functional, taxonomical and genotypic analysis into a CSR framework

We combined all the evidence gathered in this study into a schematic drawing to conceptualize the CSR life-history strategies adopted by the bacterial communities (Fig. 6). This framework aims to provide ecological guidance to understand the observed changes in community structure, genotypes and function with varying levels of disturbance, by incorporating a trait-based analysis on a whole community level.

**Fig. 6.**
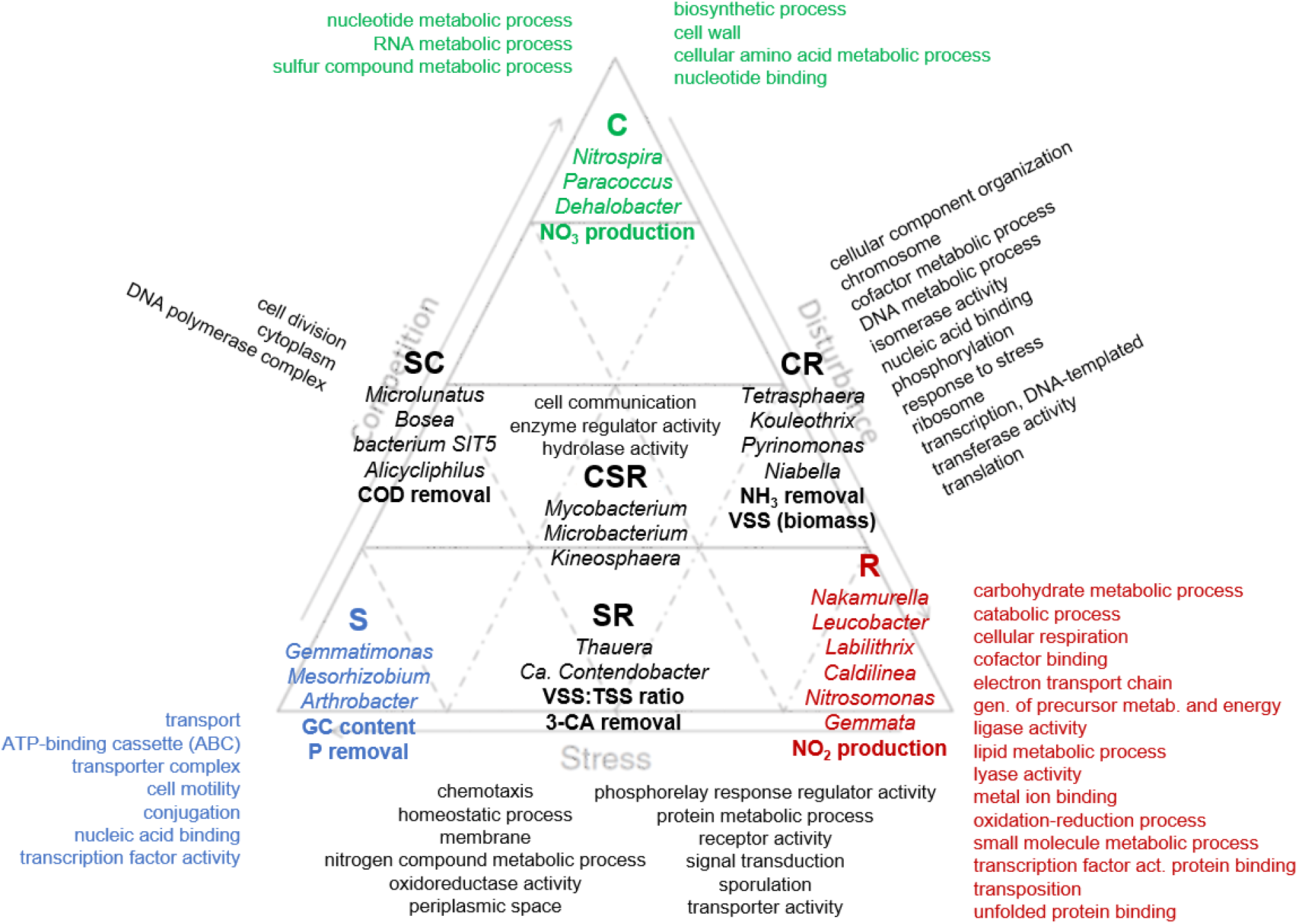
Conceptual distribution of trait complexes from IP2G database, community-level function (bold), and top 25 metagenomics genera (italicized) within the CSR life-history strategies framework. C, competitors (green); R, ruderals (red); S, stress-tolerants (blue). Intermediate strategies (CR, SR, SC, CSR) in black. The intent is to provide an interpretation of ecological drivers of changes in bacterial community structure, functional potential, and ecosystem function after succession under different levels of disturbance.

Competitors: The undisturbed reactors (L0) harboured a community with little variability across replicates and taxa well adapted to the prevailing conditions. The theoretical high efficiency associated with this life strategy (Grime and Pierce 2012) was reflected by the best NH_3_ and COD removal rates, the latter also having been highlighted as a C-trait (Chagnon *et al*. 2013), with a high biomass in comparison to the other disturbance levels. In terms of genotype, such efficiency was supported by an increase in abundance of genes associated with metabolic processes (biosynthesis, amino acids, nucleotides, RNA, sulfur compounds) and the cell wall. *Nitrospira* was the dominant nitrifier, possibly carrying the complete nitrification process (Daims *et al*. 2015) as we found NO_3_-N without NO_2_-N in the effluent for these reactors. The high abundance of *Paracoccus* could be related to denitrification processes, given the high NO_3_ availability. The dominance of some organisms under undisturbed conditions is an indication of strong competition (Hodgson *et al*. 1999).

Ruderals: Intermediately disturbed levels (L1-6) showed how disturbance prevented competitive advantages, thus allowing less prevalent seed-bank species to grow. However, variability across replicate reactors within the same disturbance levels increased for L1 to L6. This was reflected by higher standard deviations for ecosystem functions (Table S1) and dissimilarity in composition (Fig. 2, Fig. S3). Disturbance likely promoted a higher growth rate at low disturbance frequencies, since the highest biomass value was found in L1 reactors. The inefficient nutrient uptake of ruderals (Grime and Pierce 2012) was evident in the lower COD removal and partial nitrification (NO_2_ only) across some intermediately disturbed reactors. Furthermore, genes associated with metabolic processes (carbohydrates, lipids, and small molecules), as well as energy generation (electron transport chain, cellular respiration, and oxidation-reduction processes) were favoured by ruderals, which is in agreement with the notion that disturbance disrupts competition for organisms in the seed-bank (Connell 1978). Reserve material traits were suggested to be R-type (Krause *et al*. 2014), which relates to the increase in abundance of *Nakamurella*, a polysaccharide accumulator. The increase in genotypic traits of ribosome, transcription, and translation was shared among C- and R-strategists, suggesting that ribosomal activity was higher for communities at zero or low disturbance. Disturbance also promoted an increase in genes related to metabolism of aromatics and xenobiotic compounds (Fig. S6).

Stress-tolerants: At a high disturbance frequency organisms are expected to perform more maintenance functions to survive (Krause *et al*. 2014). Such tradeoffs between growth and survival might explain the low biomass value in the press-disturbed reactors (L7). Despite being slow growers and low biomass producers (Grime and Pierce 2012), S-strategists should have an efficient uptake of nutrients, which we observed as high COD and P removal values. The highest GC content was also found for this group, which might be related to higher stability as it was shown that the environment has an effect on this trait (Barberan *et al*. 2012, Foerstner *et al*. 2005). Resistance to abiotic stressors (Chagnon *et al*. 2013) as well as maintenance, membrane chemistry and uptake systems (Krause *et al*. 2014) were suggested as traits enriched in stress-tolerants. We also found increased prevalence of ATP-binding cassette transporter genes, which is reasonable as transporters are known to be vital for cell survival by counteracting undesirable changes in the cell. Sporulation was expected to increase according to Krause *et al*. (2014), but it was marked as a ruderal trait for fungi by Chagnon *et al*. (2013). We observed an increase in sporulation genes for both ruderals and stress-tolerants, as well as in genes for chemotaxis and cell motility (proposed R-traits (Krause *et al*. 2014)), suggesting that these are SR-related traits more than purely R- or S-types.

However, if communities at the maximum disturbance level of our study (L7) are identified as stress-tolerants under the CSR framework, then L7 has to be a condition of high-stress rather than high-disturbance. Grime (1977) related disturbance with biomass destruction and stress with biomass restriction. We did observe the lowest biomass for L7 reactors, but also the highest biomass for disturbed L1 ones. Thus, one could question the ability to relate our findings as CSR life-history strategies by only focusing on disturbance. How disturbance and stress are defined has been, and continues to be, debated as there is recognized inconsistency and ambiguity across studies in ecology (Borics *et al*. 2013, Rykiel 1985) and microbial ecology (Plante 2017, Sousa 1984). Our study employed varying 3-CA frequencies across disturbance levels, and higher frequencies also implied a higher capability for communities to adapt to it. Reactors at L7 experienced a continuous long-term effect of 3-CA toxicity from the beginning of the experiment, a condition known as a press disturbance (Shade *et al*. 2012a). However, it was suggested that frequency is actually the key to differentiating stress from disturbance (Borics *et al*. 2013). There is stress if events are so frequent that they prevent community structure and/or function from returning to similar pre-event dynamics, and rather shift the system toward a new course. Thus, the press-disturbed L7 communities are actually under 3-CA stress, which explains why they fit the stress-tolerant category of the CSR framework.

### Life-history strategies as valuable guidance, but not a one-size-fits-all solution

Ecological theory can provide tools to classify and interpret our observations so as to make testable predictions (Prosser *et al*. 2007). We showed here that the CSR analogy can be applied to understand ecological aspects of changes in bacterial sludge communities under varying disturbances. A separate analysis of the same experiment showed that both stochastic and deterministic assembly mechanisms were important depending on the extent of disturbance frequency, and a hump-backed β—diversity pattern was observed (Santillan *et al*. 2019). It is not surprising to find a range of mechanisms driving community succession and ecosystem responses to disturbance, as it has been shown in ecology that many post-disturbance theories can apply simultaneously to the same system (Pulsford *et al*. 2016). Their application can be intensified in microbial systems featuring characteristics (i.e. wide metabolic potential, short turnover times, genetic material transfer) that allow for the convergence of ecological and evolutionary mechanisms (Hanson *et al*. 2012). Indeed, it has been suggested that microbial community response is affected by multiple mechanisms acting concurrently (Ho *et al*. 2017).

Our proposed scheme for this study constitutes a semi-quantitative classification of life-history strategies (Fig. 6), based on results obtained through a combination of techniques. A quantitative method to allocate plant functional types in a CSR ordination triangle has been developed for grasses and other herbaceous species (Hodgson *et al*. 1999), which was tested in a study of vegetation succession in a glacier foreland (Caccianiga *et al*. 2006). The development of a similar method of CSR ordination for microorganisms could be the focus of further research efforts, but care should be taken to avoid cursory classification (Wilson and Lee 2000). More effort should be put into identifying the most relevant traits driving microbial life-history strategies (Chagnon *et al*. 2013), thus a taxonomically oriented classification within a CSR ordination (as is done for plants) may not always be relevant for microbial systems. The focus should be on reducing the vast taxonomic and functional microbial complexity, and on understanding the mechanisms driving the changes observed in ecosystem function.

### Concluding remarks

Life-history strategies frameworks enable the simplification of complex trait information into a few ecologically relevant elements, and at the same time offer a suitable management tool for characterizing changes in community structure and ecosystem function in response to perturbations (Grime 2013, Pecuchet *et al*. 2017). The results from this work are relevant for microbial ecology and represent the first time that CSR life-history strategies are: (i) proposed at the whole-community level by assessing CATs; (ii) supported by a combination of ordination methods, network analysis, and genotypic information from metagenomics and 16S rRNA gene amplicon sequencing; and (iii) evaluated for microbial communities in an environment other than soil. We contend that three-way life-history strategies using CATs can indeed reduce system complexity while offering a qualitative basis for managing microbial communities in the face of inherent uncertainty. Ultimately, they constitute a valuable and suitable tool for the exploration of microbial community changes upon disturbance, with the potential to advance the search for the mechanisms operating on a community level.

## Experimental Procedures

### Experimental design and functional parameters

A microcosm experimental setup was operated through time using different levels of 3-CA disturbance as described by Santillan *et al*. (2019). Briefly, sequencing batch bioreactors (20-mL working volume) were inoculated with activated sludge from a full-scale plant in Singapore and operated for 35 days. The daily complex synthetic feed targeted concentrations of 590 (±15.4) mg COD L^-1^ and 92 (±2.5) mg N L^-1^ in mixed liquor, and included 3-CA (70 mg L^-1^) at varying frequencies. Whenever 3-CA was added, the C- and N-containing compounds in the feed were proportionally reduced to maintain the above COD and N target concentrations. Phosphates (357 ±8.4 mg P L^-1^) were used to buffer the medium and maintain a pH of around 7.5 to facilitate the nitrification process. Eight levels of disturbance were set in triplicate independent reactors (n = 24), which received 3-CA either never (undisturbed, L0), every 7, 6, 5, 4, 3, or 2 days (intermediately-disturbed, L1-6), or every day (press-disturbed, L7). Community-level function, in the form of process performance parameters, was measured weekly in accordance with Standard Methods (APHA-AWWA-WEF 2005) where appropriate, and targeted soluble chemical oxygen demand (COD), nitrogen species (ammonium, nitrite, and nitrate ions), phosphorus (phosphate ions), volatile (VSS) and total suspended solids (TSS), and 3-CA. Sludge samples (2 mL) were collected initially (d0) and weekly from d14 onwards for DNA extraction. Temporal dynamics of ecosystem function are detailed in Santillan *et al*. (2019), while only functional data from d35 were employed for this study.

### 16S rRNA amplicon sequencing and reads processing

Bacterial 16S rRNA amplicon sequencing was done at the SCELSE sequencing facility in two steps (for details see Supporting Methods). Primer set 341f/785r targeted the V3-V4 variable regions of the 16S rRNA gene (Thijs *et al*. 2017). The libraries were sequenced on an Illumina MiSeq platform (v.3). Sequenced sample libraries were processed following the DADA2 bioinformatics pipeline (Callahan *et al*. 2016) using the dada2 R-package (v.1.3.3). DADA2 allows inference of exact amplicon sequence variants (ASVs) providing several benefits over traditional OTU clustering methods (Callahan *et al*. 2017). Illumina sequencing adaptors and PCR primers were trimmed prior to quality filtering. Sequences were truncated after 280 and 255 nucleotides for forward and reverse reads, respectively, the length at which average quality dropped below a Phred score of 20. After truncation, reads with expected error rates higher than 3 and 5 for forward and reverse reads were removed. After filtering, error rate learning, ASV inference and denoising, reads were merged with a minimum overlap of 20 bp. Chimeric sequences (0.07% on average) were identified and removed. For a total of 99 samples, 12291 reads were kept on average per sample after processing, representing 13.2% of the average input reads. Taxonomy was assigned using the SILVA database (v.132) (Glöckner *et al*. 2017).

### Metagenomics sequencing and reads processing

Metagenomics library preparation and sequencing were as described by Santillan *et al*. (2019). Briefly, libraries were sequenced in one lane on an Illumina HiSeq2500 sequencer in rapid mode at a read-length of 250 bp. Taxonomic assignment of metagenomics reads was done following the method described by Ilott *et al*. (2016). The lowest common ancestor approach implemented in MEGAN (Community Edition v.6.5.5 (Huson *et al*. 2016)) was used to assign taxonomy to the NCBI-NR aligned reads. On average, 48.2% of the high-quality reads were assigned to cellular organisms, from which in turn 98% were assigned to the bacterial domain. Additionally, functional potential data were obtained from the metagenomics dataset using MEGAN. Analysis was done at a level of trait complex classification to capture broad changes in genotypes due to disturbance. From the four databases available in MEGAN, InterPro2Go (Finn *et al*. 2017) (IP2G) had the most hits with regards to the total bacterial reads (91.5%), and was thus employed for the main analysis. Other available databases had fewer hits: 38.4% (COG), 26.8% (SEED), and 25.9% (KEGG).

### Bacterial community and genotypic analysis

Community structure was assessed on genus-level metagenomics and 16S rRNA gene amplicon sequencing data by a combination of ordination methods and multivariate tests employing PRIMER (v.7) (Clarke and Gorley 2015). Normalized abundance data were employed, with square root transformation applied to reduce the weight of the most abundant genera. To evaluate how community function related to changes in community structure patterns across disturbance levels, constrained canonical analysis of principal coordinates (CAP) ordination including Pearson’s correlation vectors of normalized community function data were used. Permutational multivariate analysis of variance (PERMANOVA) was performed on Bray-Curtis dissimilarity matrixes (Anderson and Walsh 2013) to test whether communities at the metagenomics genus-level differed with disturbance. Factors were considered fixed. Homogeneity of multivariate dispersions was tested by PERMDISP (Anderson 2006). P-values were calculated using 9,999 permutations. Community changes due to disturbance were further assessed by general linear multivariate models (GLMMs) to deal with mean-variance relationships (Warton *et al*. 2012), using the *mvabund* R package (Wang *et al*. 2012) to fit the 500 most abundant metagenomics genera to a negative binomial distribution as described by Santillan *et al*. (2019). To evaluate how genotypes related with the observed differences in bacterial community structure, distance-based redundancy analysis (dbRDA) (McArdle and Anderson 2001) was employed using IP2G trait-complexes as predictor variables. Pearson’s correlation vectors (r > 0.20) were overlaid on the dbRDA plot.

Network analysis was performed with Gephi (v.0.9.2) using Pearson’s correlation matrixes for the top 200 genera, for both metagenomics and 16S rRNA gene amplicon sequencing datasets, as well as trait complexes and the top 200 genes from the IP2G database on the metagenomics dataset. Input correlation matrixes were generated using the *Hmisc* and *corrplot* R packages. Non-significant correlations (r < 0.5, r < 0.6) were filtered out. Node clusters were defined through modularity class calculated using the Louvain method, and were coloured through identification of representative nodes at undisturbed, intermediately-disturbed and press-disturbed levels. Layout was adjusted using the Fruchterman Reingold method with default parameters. Node size was adjusted by degree, while edge thickness was adjusted by correlation strength. Changes in bacterial genera and genotypes abundances were assessed through heat maps and cluster analysis using MEGAN (Huson *et al*. 2016). Univariate Welch’s ANOVA tests were employed to evaluate the effect of disturbance on IP2G trait complexes, using IBM SPSS (v.25). All calculated P-values were adjusted for multiple comparisons using a false discovery rate (FDR) (Benjamini and Hochberg 1995) of 10%.

## Data availability

DNA sequencing data are available at NCBI BioProjects with accession number: 389377. Additional Supporting Information may be found in the online version of this article at the publisher’s web site.

### Acknowledgements

This research was supported by the Singapore National Research Foundation and Ministry of Education under the Research Centre of Excellence Program. We thank D. I. Drautz-Moses for her guidance with the sequencing pipelines employed. We acknowledge M. Holyoak and K. M. Scow for critical feedback. E.S. was partially supported by a Fulbright fellowship.

## Author Contributions

ES and SW conceived this study. ES and HS designed the experiment. SW obtained the funding for the study. ES and HS performed the experiments and laboratory analyses. ES conducted all molecular analyses except library preparation and sequencing. ES performed the 16S rRNA gene and FC the metagenomics bioinformatics analyses. ES interpreted the data, generated the results, and elaborated the main arguments in the manuscript. ES and SW wrote the manuscript. HC and FC critically reviewed the manuscript.

