## Supporting Information for "Trait-based life-history strategies explain succession scenario for complex bacterial communities under varying disturbance"

**Supporting Methods**

*16S rRNA amplicon library preparation and sequencing*

For the first PCR stage, each reaction (25 μL) contained 12.5 μL of HiFi Hotstart Readymix (Kapa Biosystems), 9.5 μL of nuclease free water, 0.5 μL (each) of forward and reverse primers (10 μM) and 2 μL of DNA template (6 ng $\mu L^{-1}$). Primer set 341f/785r targeted the V3-4 variable regions of the 16S rRNA gene (Thijs et al. 2017). Thermocycler settings were: Initial denaturation at 95°C for 2 min, 25 cycles of 95°C for 30 s, 58°C for 15 s, 72°C for 30 s, and final elongation at 72°C for 2 min. PCR reactions were all run in duplicate and pooled subsequently. Amplicon libraries were purified using the Agencourt AMpure XP bead protocol (Beckmann Coulter). Library concentration was measured with Qubit 3.0 fluorometer (Thermo Fisher Scientific) and quality validated with a Tapestation 2200 (Agilent).

The second stage PCR (Indexing PCR) was performed according to the recommendations in Illumina's '16S Metagenomic Sequencing Library Preparation' application note. This step uses a limited 8-cycles PCR to complete the Illumina sequencing adapters and add dual-index barcodes to the amplicon target. Five microliters of the intermediate PCR product from the first stage were used as template for the indexing PCR and samples were amplified with 8 PCR cycles. Nextera XT v2 indices were used for dual-index barcoding to allow pooling of the amplicon targets for sequencing.

Finished amplicon libraries were quantitated using QuantiFluor dsDNA assay (Promega) and the average library size was determined on a Tapestation 4200 (Agilent). Library concentrations were then normalized to 4nM and validated by qPCR on a QuantStudio-3 system (Applied Biosystems), using the Kapa library quantification kit for Illumina platforms (Kapa Biosystems). The libraries were then pooled at equimolar concentrations and sequenced on an Illumina MiSeq platform (v.3) with 20% PhiX spike-in and at a read-length of 300bp paired-end. Sequencing was done at SCELSE's core sequencing facility.

### Supporting Figures

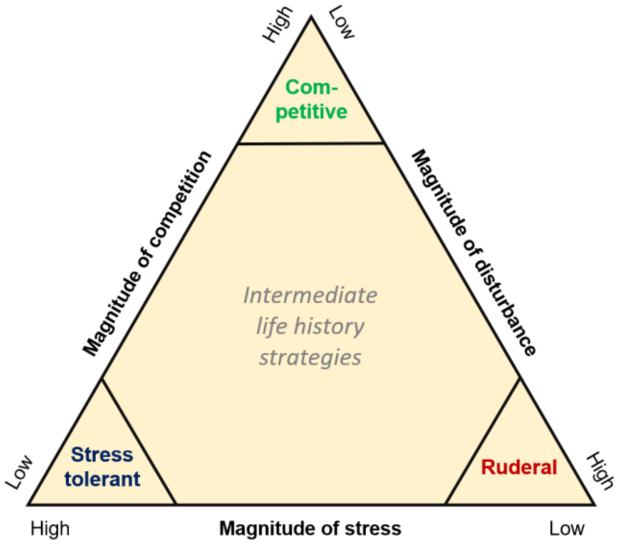


**Fig. S1.** Schematic representation of the CSR framework for plant communities (adapted from Grime 1977).

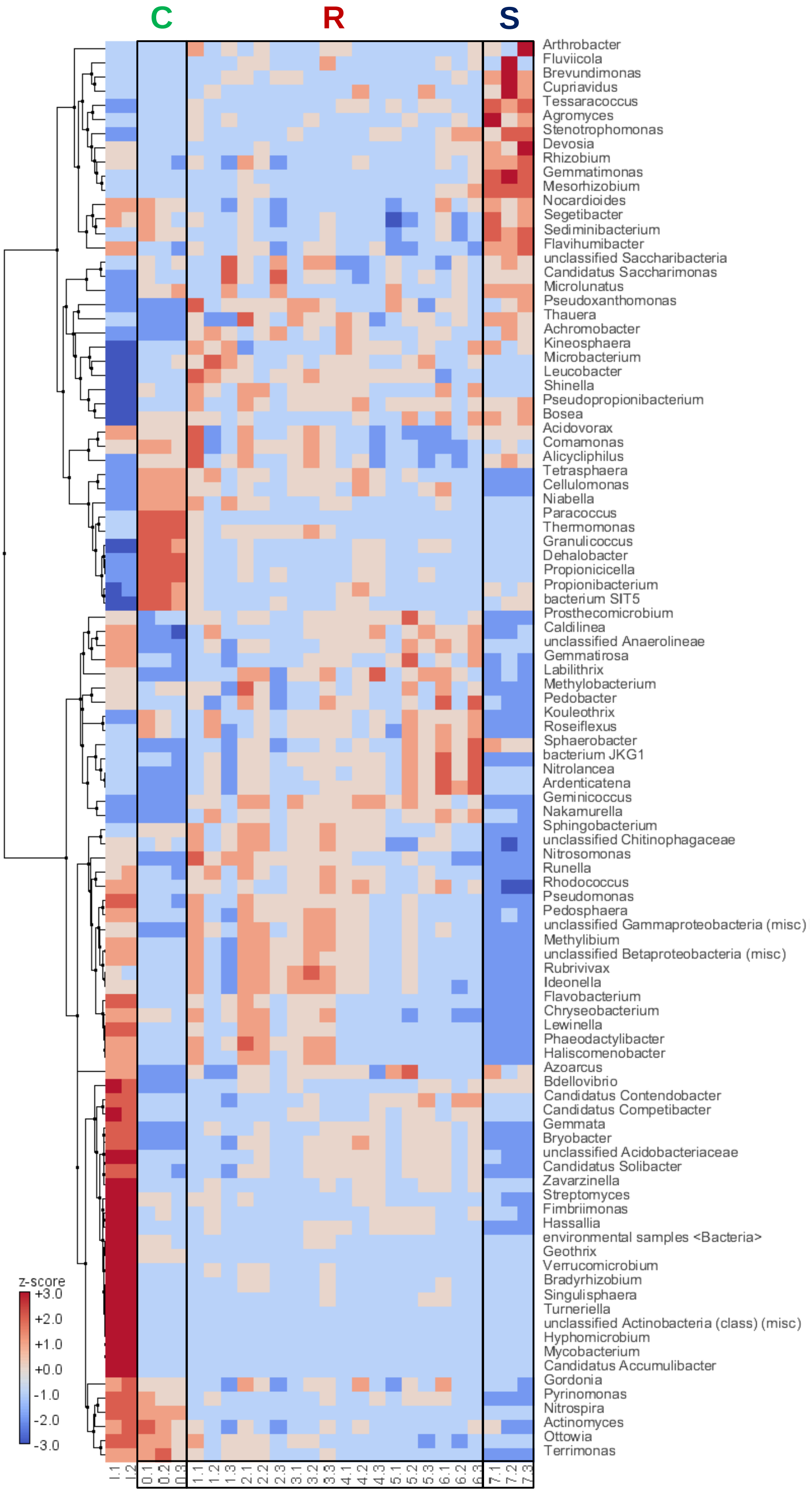


**Fig. S2.** Clustered heat map for the 100 most abundant genera across reactors. First two columns (I.1, I.2) are replicates from the WWTP inoculum (d0). The subsequent column legends represents disturbance level and replicate reactors on day 35 (n = 24). Rectangles highlight taxa groups prevailing at different CSR life-history strategies at the community-level: C, competitors (L0); R, ruderals (L1-6); S, stress-tolerants (L7).

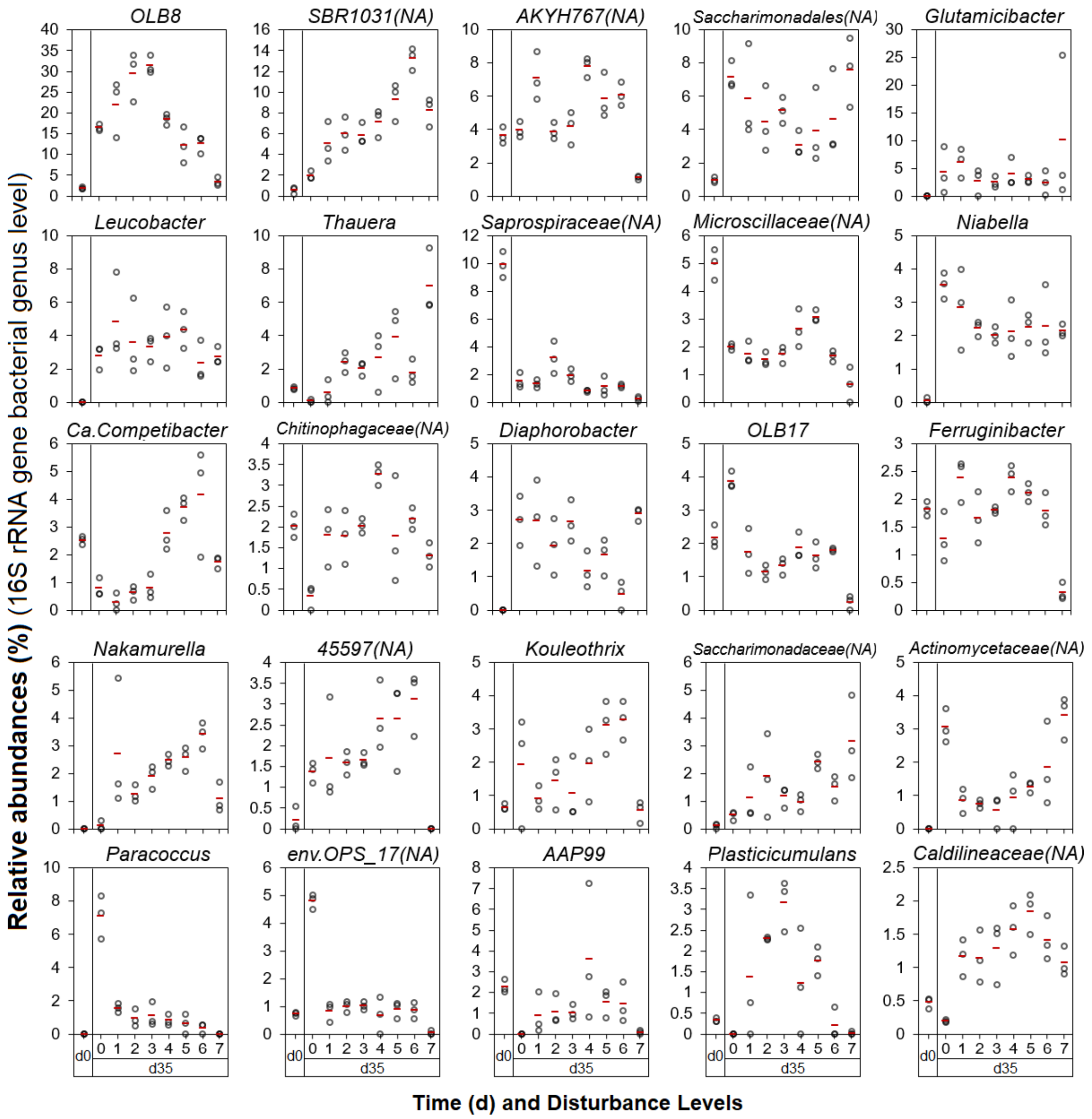


**Fig. S3.** Relative abundance comparisons to discern dominant taxa at different disturbance levels across reactors (n = 24). Top 25 genus-level taxa assessed through 16S rRNA gene amplicon sequencing at d35. Relative abundances at d0 included for comparison with initial conditions. Open circles indicate replicate values, while red lines display the average for a particular level. Note that *Ca.* Competibacter and *Glutamicibacter* are named *Ca.* Contendobacter and *Arthrobacter*, respectively for the metagenomics genus-level dataset. NA: not assigned at genus level (last taxonomic category assigned shown).

| d | 0 | 0 | 0 |
|---|---|---|---|
| L | I | I | I |
| **Genera** | **I.1** | **I.2** | **I.3** |
| Saprospiraceae(NA) | 11 | 9.8 | 9.0 |
| Chitinophagales(NA) | 5.2 | 5.7 | 5.1 |
| Microscillaceae(NA) | 5.5 | 5.1 | 4.4 |
| AKYH767(NA) | 3.5 | 4.2 | 3.2 |
| Nitrospira | 3.4 | 3.8 | 3.3 |
| Mycobacterium | 2.9 | 2.5 | 2.1 |
| Ca. Competibacter | 2.4 | 2.7 | 2.5 |
| Rhodanobacteraceae(NA) | 2.4 | 2.6 | 2.6 |
| Rhodocyclaceae(NA) | 2.5 | 2.5 | 2.0 |
| env.OPS_17(NA) | 2.2 | 2.6 | 2.0 |
| OLB17 | 2.1 | 2.6 | 1.9 |
| Chitinophagaceae(NA) | 2.0 | 2.3 | 1.8 |
| IMCC26207 | 2.0 | 2.3 | 1.7 |
| OLB8 | 1.8 | 2.3 | 1.8 |
| Ferruginibacter | 2.0 | 1.8 | 1.7 |
| 67-14(NA) | 1.8 | 1.6 | 1.0 |
| Azospira | 1.4 | 1.5 | 1.4 |
| Novosphingobium | 1.3 | 1.3 | 1.6 |
| Ottowia | 1 | 1.2 | 1.4 |
| Planctomycetales(NA) | 1.4 | 1.4 | 1.1 |

| % |
|---|
| 0 |
| 1 |
| 2 |
| 3 |
| 4 |
| 5 |
| 10 |
| 15 |
| 20 |
| 25 |
| 30 |
| 40 |

| d | 0 | 0 | 0 | 14 | 14 | 14 | 21 | 21 | 21 | 28 | 28 | 28 | 35 | 35 | 35 |
|---|---|---|---|---|---|---|---|---|---|---|---|---|---|---|---|
| L | I | I | I | 0 | 0 | 0 | 0 | 0 | 0 | 0 | 0 | 0 | 0 | 0 | 0 |
| **Genera** | **I.1** | **I.2** | **I.3** | **0.1** | **0.2** | **0.3** | **0.1** | **0.2** | **0.3** | **0.1** | **0.2** | **0.3** | **0.1** | **0.2** | **0.3** |
| OLB8 | 1.8 | 2.3 | 1.8 | 14 | 15 | 12 | 22 | 23 | 23 | 25 | 28 | 29 | 16 | 17 | 16 |
| Saccharimonadales(NA) | 1.1 | 0.8 | 0.9 | 16 | 11 | 14 | 17 | 12 | 14 | 7.7 | 7.6 | 4.6 | 8.1 | 6.7 | 6.6 |
| Paracoccus | 0.0 | 0.0 | 0.0 | 2.0 | 4.2 | 1.6 | 3.8 | 5.1 | 4.1 | 4.0 | 6.6 | 4.8 | 5.7 | 7.3 | 8.3 |
| AAP99 | 0.8 | 0.8 | 0.7 | 0.0 | 0.6 | 0.6 | 1.1 | 1.1 | 1.0 | 2.9 | 3.2 | 3.4 | 4.9 | 5.0 | 4.5 |
| Glutamicibacter | 0.0 | 0.0 | 0.0 | 3.9 | 2.7 | 1.4 | 7.5 | 5.0 | 2.1 | 7.5 | 0.9 | 2.5 | 8.9 | 0.7 | 3.3 |
| AKYH767(NA) | 3.5 | 4.2 | 3.2 | 4.3 | 4.8 | 6.4 | 3.9 | 4.7 | 6.6 | 3.5 | 2.8 | 4.3 | 3.5 | 3.8 | 4.5 |
| OLB17 | 2.1 | 2.6 | 1.9 | 1.6 | 1.7 | 1.4 | 1.5 | 0.7 | 1.8 | 2.6 | 2.9 | 1.7 | 3.7 | 4.2 | 3.7 |
| Niabella | 0.0 | 0.0 | 0.1 | 4.0 | 2.5 | 3.1 | 3.1 | 1.9 | 2.8 | 2.3 | 2.4 | 2.7 | 3.1 | 3.9 | 3.6 |
| Actinomycetaceae(NA) | 0.0 | 0.0 | 0.0 | 0.8 | 0.7 | 0.0 | 1.6 | 1.7 | 2.3 | 2.9 | 1.8 | 3.4 | 2.6 | 3.0 | 3.6 |
| Leucobacter | 0.0 | 0.0 | 0.0 | 4.7 | 7.0 | 7.2 | 3.6 | 7.8 | 4.8 | 1.8 | 2.7 | 3.9 | 1.9 | 3.2 | 3.2 |
| Diaphorobacter | 0.0 | 0.0 | 0.0 | 2.0 | 2.1 | 2.2 | 2.1 | 3.0 | 3.4 | 1.9 | 1.7 | 3.6 | 2.0 | 2.7 | 3.4 |
| Micropruina | 0.0 | 0.0 | 0.0 | 2.2 | 2.0 | 1.5 | 3.2 | 4.7 | 3.2 | 1.0 | 1.5 | 4.4 | 3.3 | 2.5 | 2.3 |
| Nitrospira | 3.4 | 3.8 | 3.3 | 2.7 | 3.0 | 2.8 | 1.9 | 1.9 | 1.7 | 3.3 | 1.9 | 2.9 | 2.4 | 2.8 | 2.7 |
| Microscillaceae(NA) | 5.5 | 5.1 | 4.4 | 1.3 | 1.2 | 1.4 | 1.2 | 0.5 | 1.3 | 1.6 | 1.3 | 1.5 | 2.0 | 1.9 | 2.1 |
| SBR1031(NA) | 0.1 | 0.8 | 0.7 | 1.7 | 1.8 | 2.4 | 2.2 | 1.6 | 1.1 | 2.5 | 3.3 | 0.6 | 1.7 | 2.5 | 1.7 |
| Kouleothrix | 0.6 | 0.6 | 0.8 | 1.1 | 0.9 | 2.7 | 2.8 | 0.6 | 2.9 | 3.9 | 3.7 | 0.9 | 2.6 | 3.2 | 0.0 |
| Thermomonas | 0.0 | 0.0 | 0.0 | 1.9 | 1.4 | 2.3 | 0.5 | 0.7 | 0.9 | 1.0 | 1.3 | 1.4 | 1.5 | 1.6 | 2.2 |
| Bosea | 0.0 | 0.0 | 0.0 | 1.1 | 0.5 | 1.0 | 1.2 | 1.2 | 1.3 | 0.8 | 1.4 | 1.4 | 1.1 | 1.8 | 1.7 |
| Saprospiraceae(NA) | 11 | 9.8 | 9.0 | 7.2 | 5.9 | 7.2 | 3.4 | 3.7 | 4.8 | 3.0 | 3.6 | 3.0 | 1.3 | 2.1 | 1.1 |
| 45597(NA) | 0.1 | 0.5 | 0.0 | 0.0 | 0.3 | 0.1 | 0.4 | 0.4 | 0.0 | 0.9 | 0.8 | 0.9 | 1.4 | 1.6 | 1.1 |

| d | 0 | 0 | 0 | 14 | 14 | 14 | 21 | 21 | 21 | 28 | 28 | 28 | 35 | 35 | 35 |
|---|---|---|---|---|---|---|---|---|---|---|---|---|---|---|---|
| L | I | I | I | 1 | 1 | 1 | 1 | 1 | 1 | 1 | 1 | 1 | 1 | 1 | 1 |
| **Genera** | **I.1** | **I.2** | **I.3** | **1.1** | **1.2** | **1.3** | **1.1** | **1.2** | **1.3** | **1.1** | **1.2** | **1.3** | **1.1** | **1.2** | **1.3** |
| OLB8 | 1.8 | 2.3 | 1.8 | 38 | 21 | 33 | 25 | 34 | 27 | 29 | 31 | 27 | 27 | 14 | 25 |
| AKYH767(NA) | 3.5 | 4.2 | 3.2 | 9 | 9 | 5 | 6 | 5 | 9 | 6.6 | 4.6 | 7.9 | 5.8 | 6.8 | 8.7 |
| Glutamicibacter | 0.0 | 0.0 | 0.0 | 1.4 | 1.1 | 3.1 | 2.5 | 1.2 | 2.1 | 5.7 | 2.2 | 4.7 | 6.7 | 3.3 | 8.4 |
| Saccharimonadales(NA) | 1.1 | 0.8 | 0.9 | 2.6 | 7.7 | 3.1 | 6.7 | 3.5 | 5.2 | 6.0 | 4.7 | 7.3 | 4.0 | 4.4 | 9.2 |
| SBR1031(NA) | 0.1 | 0.8 | 0.7 | 3.5 | 2.9 | 6.7 | 4.9 | 4.2 | 4.7 | 4.1 | 5.5 | 3.5 | 4.5 | 7.2 | 3.4 |
| Leucobacter | 0.0 | 0.0 | 0.0 | 1.6 | 7.5 | 3.0 | 4.1 | 5.0 | 3.3 | 3.7 | 4.9 | 3.4 | 3.5 | 7.8 | 3.2 |
| Niabella | 0.0 | 0.0 | 0.1 | 1.2 | 1.1 | 1.6 | 2.3 | 1.7 | 2.1 | 2.0 | 1.7 | 2.6 | 3.0 | 1.6 | 4.0 |
| Nakamurella | 0.0 | 0.0 | 0.0 | 0.0 | 0.0 | 0.1 | 0.5 | 0.8 | 0.0 | 0.6 | 1.4 | 0.8 | 1.6 | 5.4 | 1.1 |
| Diaphorobacter | 0.0 | 0.0 | 0.0 | 1.7 | 1.1 | 2.6 | 2.8 | 1.1 | 3.1 | 2.8 | 1.1 | 1.8 | 3.9 | 1.3 | 2.8 |
| Ferruginibacter | 2.0 | 1.8 | 1.7 | 1.9 | 2.0 | 1.5 | 2.5 | 2.3 | 2.9 | 2.5 | 2.4 | 2.9 | 1.9 | 2.6 | 2.6 |
| Chitinophagaceae(NA) | 2.0 | 2.3 | 1.8 | 0.0 | 0.0 | 0.4 | 0.0 | 0.4 | 0.1 | 0.6 | 0.5 | 0.3 | 1.9 | 2.4 | 1.0 |
| OLB17 | 2.1 | 2.6 | 1.9 | 1.9 | 1.4 | 1.4 | 1.8 | 1.3 | 2.6 | 1.1 | 1.4 | 1.6 | 1.1 | 1.7 | 2.5 |
| Microscillaceae(NA) | 5.5 | 5.1 | 4.4 | 1.3 | 1.8 | 0.3 | 3.0 | 2.5 | 3.0 | 2.2 | 2.5 | 2.9 | 1.5 | 2.2 | 1.5 |
| 45597(NA) | 0.1 | 0.5 | 0.0 | 0.2 | 0.2 | 0.7 | 0.7 | 1.0 | 0.6 | 1.0 | 1.4 | 0.9 | 1.0 | 3.2 | 0.9 |
| Paracoccus | 0.0 | 0.0 | 0.0 | 0.0 | 0.9 | 0.0 | 1.3 | 0.9 | 0.8 | 1.4 | 1.0 | 1.0 | 1.8 | 1.3 | 1.6 |
| Plasticicumulans | 0.4 | 0.3 | 0.3 | 2.4 | 1.7 | 0.3 | 1.0 | 0.2 | 0.3 | 0.2 | 0.3 | 0.4 | 3.4 | 0.0 | 0.7 |
| Saprospiraceae(NA) | 11 | 9.8 | 9.0 | 12 | 7.0 | 9.2 | 3.5 | 5.3 | 2.7 | 2.9 | 3.4 | 1.7 | 1.6 | 1.3 | 1.0 |
| PLTA13(NA) | 0.6 | 0.6 | 0.9 | 0.6 | 0.5 | 0.6 | 0.9 | 1.1 | 0.9 | 1.1 | 1.2 | 1.4 | 0.6 | 2.0 | 1.1 |
| Caldilineaceae(NA) | 1 | 0.4 | 0.5 | 1.5 | 1.6 | 1.5 | 1.7 | 1.4 | 2.5 | 2.3 | 2.1 | 1.8 | 1.2 | 1.4 | 0.9 |
| Saccharimonadaceae(NA) | 0.0 | 0.1 | 0.2 | 0.0 | 0.6 | 0.6 | 0.5 | 0.2 | 0.2 | 0.8 | 0.9 | 1.3 | 0.6 | 0.6 | 2.2 |

| d | 0 | 0 | 0 | 14 | 14 | 14 | 21 | 21 | 21 | 28 | 28 | 28 | 35 | 35 | 35 |
|---|---|---|---|---|---|---|---|---|---|---|---|---|---|---|---|
| L | I | I | I | 2 | 2 | 2 | 2 | 2 | 2 | 2 | 2 | 2 | 2 | 2 | 2 |
| **Genera** | **I.1** | **I.2** | **I.3** | **2.1** | **2.2** | **2.3** | **2.1** | **2.2** | **2.3** | **2.1** | **2.2** | **2.3** | **2.1** | **2.2** | **2.3** |
| OLB8 | 1.8 | 2.3 | 1.8 | 25 | 25 | 20 | 42 | 37 | 36 | 31 | 32 | 22 | 34 | 32 | 23 |
| SBR1031(NA) | 0.1 | 0.8 | 0.7 | 5 | 4 | 4 | 3 | 3 | 5 | 6.7 | 6.9 | 7.1 | 4.4 | 7.6 | 5.9 |
| Saccharimonadales(NA) | 1.1 | 0.8 | 0.9 | 11 | 7 | 11 | 2.7 | 4.0 | 5.2 | 5.3 | 3.7 | 5.6 | 3.9 | 2.8 | 6.6 |
| AKYH767(NA) | 3.5 | 4.2 | 3.2 | 7.2 | 7.5 | 7.8 | 4.3 | 6.0 | 5.5 | 3.7 | 4.0 | 4.7 | 3.8 | 3.5 | 4.4 |
| Leucobacter | 0.0 | 0.0 | 0.0 | 7.5 | 6.5 | 7.7 | 2.2 | 2.2 | 4.1 | 2.0 | 1.6 | 5.2 | 2.6 | 1.9 | 6.2 |
| Saprospiraceae(NA) | 11 | 9.8 | 9.0 | 7.9 | 7.2 | 9.8 | 7.4 | 5.2 | 8.6 | 5.0 | 3.6 | 2.5 | 4.4 | 3.1 | 2.1 |
| Glutamicibacter | 0.0 | 0.0 | 0.0 | 0.5 | 1.5 | 1.5 | 1.2 | 1.7 | 0.8 | 0.0 | 3.0 | 2.2 | 0.0 | 4.5 | 3.7 |
| Thauera | 0.9 | 0.8 | 0.8 | 0.0 | 0.0 | 0.4 | 0.6 | 0.4 | 0.7 | 0.9 | 0.7 | 1.0 | 3.0 | 1.8 | 2.5 |
| Plasticicumulans | 0.4 | 0.3 | 0.3 | 0.0 | 0.2 | 1.1 | 1.0 | 0.9 | 0.5 | 2.7 | 3.5 | 1.4 | 2.3 | 2.3 | 2.3 |
| Niabella | 0.0 | 0.0 | 0.1 | 1.4 | 1.5 | 1.3 | 2.5 | 2.8 | 1.4 | 2.0 | 1.9 | 1.6 | 2.4 | 2.0 | 2.3 |
| Diaphorobacter | 0.0 | 0.0 | 0.0 | 3.3 | 3.9 | 3.8 | 1.1 | 1.2 | 1.8 | 1.0 | 1.8 | 1.3 | 2.7 | 1.1 | 2.0 |
| Saccharimonadaceae(NA) | 0.0 | 0.1 | 0.2 | 0.2 | 0.3 | 0.3 | 0.3 | 0.0 | 0.3 | 1.1 | 0.0 | 1.5 | 1.8 | 0.4 | 3.4 |
| Chitinophagaceae(NA) | 2.0 | 2.3 | 1.8 | 0.0 | 0.0 | 0.0 | 0.1 | 0.3 | 0.0 | 0.8 | 0.3 | 0.4 | 2.4 | 1.8 | 1.1 |
| Ferruginibacter | 2.0 | 1.8 | 1.7 | 1.6 | 1.8 | 1.8 | 2.0 | 2.5 | 2.2 | 2.0 | 2.0 | 2.4 | 1.6 | 1.2 | 2.1 |
| 45597(NA) | 0.1 | 0.5 | 0.0 | 0.2 | 0.3 | 0.4 | 1.1 | 0.8 | 0.8 | 1.2 | 1.4 | 1.4 | 1.9 | 1.6 | 1.3 |
| Microscillaceae(NA) | 5.5 | 5.1 | 4.4 | 1.5 | 1.6 | 1.8 | 2.0 | 2.9 | 2.7 | 2.2 | 1.9 | 3.1 | 1.4 | 1.4 | 1.8 |
| Kouleothrix | 1 | 0.6 | 0.8 | 1 | 1.6 | 0.9 | 0.4 | 1.8 | 2.2 | 2.5 | 2.3 | 2.2 | 1.7 | 0.6 | 2.1 |
| OPB56(NA) | 0.0 | 0.0 | 0.1 | 0.0 | 0.0 | 0.0 | 0.2 | 0.2 | 0.0 | 0.9 | 0.7 | 0.1 | 1.6 | 2.2 | 0.5 |
| Burkholderiaceae(NA) | 1 | 1.1 | 1.0 | 0.0 | 0.1 | 0.0 | 0.0 | 0.2 | 0.0 | 0.5 | 0.0 | 0.0 | 1.9 | 1.2 | 1.1 |
| Nakamurella | 0.0 | 0.0 | 0.0 | 0.3 | 0.2 | 0.0 | 0.0 | 0.4 | 0.4 | 0.9 | 0.9 | 0.9 | 1.0 | 1.6 | 1.2 |

| d | 0 | 0 | 0 | 14 | 14 | 14 | 21 | 21 | 21 | 28 | 28 | 28 | 35 | 35 | 35 |
|---|---|---|---|---|---|---|---|---|---|---|---|---|---|---|---|
| L | I | I | I | 3 | 3 | 3 | 3 | 3 | 3 | 3 | 3 | 3 | 3 | 3 | 3 |
| **Genera** | **I.1** | **I.2** | **I.3** | **3.1** | **3.2** | **3.3** | **3.1** | **3.2** | **3.3** | **3.1** | **3.2** | **3.3** | **3.1** | **3.2** | **3.3** |
| OLB8 | 1.8 | 2.3 | 1.8 | 27 | 22 | 28 | 34 | 31 | 31 | 25 | 25 | 29 | 30 | 34 | 30 |
| SBR1031(NA) | 0.1 | 0.8 | 0.7 | 2 | 4 | 3 | 6 | 6 | 6 | 6.6 | 6.3 | 5.6 | 5.2 | 5.2 | 7.1 |
| Saccharimonadales(NA) | 1.1 | 0.8 | 0.9 | 8.0 | 11 | 9.3 | 3.3 | 5.2 | 4.4 | 2.3 | 7.4 | 6.0 | 6.0 | 5.1 | 4.4 |
| AKYH767(NA) | 3.5 | 4.2 | 3.2 | 7.4 | 7.5 | 8.5 | 4.8 | 5.1 | 4.6 | 4.1 | 4.0 | 4.6 | 4.4 | 3.1 | 5.0 |
| Leucobacter | 0.0 | 0.0 | 0.0 | 5.0 | 7.4 | 3.9 | 1.6 | 3.7 | 2.4 | 2.6 | 3.6 | 2.4 | 2.5 | 3.7 | 3.8 |
| Plasticicumulans | 0.4 | 0.3 | 0.3 | 1.1 | 1.4 | 0.7 | 0.4 | 0.5 | 0.3 | 1.1 | 0.7 | 2.1 | 2.5 | 3.4 | 3.6 |
| Diaphorobacter | 0.0 | 0.0 | 0.0 | 1.4 | 1.6 | 1.2 | 2.8 | 2.9 | 3.1 | 1.7 | 1.7 | 3.1 | 2.1 | 2.5 | 3.3 |
| Glutamicibacter | 0.0 | 0.0 | 0.0 | 7.1 | 0.8 | 0.8 | 8.6 | 0.8 | 1.8 | 8.6 | 2.4 | 3.5 | 1.6 | 2.2 | 3.7 |
| Thauera | 0.9 | 0.8 | 0.8 | 0.9 | 0.5 | 1.1 | 0.2 | 0.2 | 0.0 | 0.5 | 0.5 | 2.5 | 2.3 | 2.3 | 1.6 |
| Chitinophagaceae(NA) | 2.0 | 2.3 | 1.8 | 0.0 | 0.0 | 0.0 | 0.5 | 0.1 | 0.2 | 0.9 | 0.4 | 0.8 | 2.2 | 2.0 | 1.9 |
| Niabella | 0.0 | 0.0 | 0.1 | 1.3 | 1.2 | 1.1 | 1.5 | 1.7 | 1.7 | 1.5 | 1.4 | 2.3 | 2.0 | 1.8 | 2.3 |
| Saprospiraceae(NA) | 11 | 9.8 | 9.0 | 7.1 | 7.9 | 8.2 | 5.4 | 4.9 | 5.6 | 2.0 | 2.2 | 3.5 | 2.4 | 1.9 | 1.5 |
| Nakamurella | 0.0 | 0.0 | 0.0 | 0.0 | 0.3 | 0.0 | 0.5 | 0.5 | 0.5 | 1.8 | 0.3 | 1.0 | 2.2 | 2.0 | 1.4 |
| Ferruginibacter | 2.0 | 1.8 | 1.7 | 2.0 | 1.9 | 1.7 | 1.8 | 2.0 | 2.8 | 1.9 | 2.2 | 2.4 | 1.7 | 1.8 | 1.9 |
| Microscillaceae(NA) | 5.5 | 5.1 | 4.4 | 2.1 | 1.9 | 1.5 | 1.9 | 3.1 | 2.4 | 2.7 | 2.6 | 2.6 | 1.8 | 2.0 | 1.4 |
| 45597(NA) | 0.1 | 0.5 | 0.0 | 0.0 | 0.3 | 0.5 | 0.5 | 0.9 | 1.1 | 1.3 | 1.5 | 1.6 | 1.6 | 1.5 | 1.8 |
| PLTA13(NA) | 1 | 0.6 | 0.9 | 0 | 0.4 | 0.3 | 0.4 | 0.7 | 0.8 | 1.3 | 0.9 | 1.0 | 1.0 | 1.9 | 1.7 |
| OLB17 | 2.1 | 2.6 | 1.9 | 1.4 | 1.6 | 1.6 | 1.3 | 1.5 | 1.2 | 1.1 | 1.5 | 1.4 | 1.4 | 1.5 | 1.1 |
| Caldilineaceae(NA) | 1 | 0.4 | 0.5 | 1.7 | 1.5 | 2.1 | 2.3 | 2.4 | 2.4 | 1.7 | 1.8 | 1.7 | 1.5 | 0.7 | 1.6 |
| Saccharimonadaceae(NA) | 0.0 | 0.1 | 0.2 | 0.0 | 0.0 | 0.4 | 0.2 | 0.8 | 0.8 | 0.2 | 2.5 | 1.5 | 0.7 | 1.4 | 1.4 |

| d | 0 | 0 | 0 | 14 | 14 | 14 | 21 | 21 | 21 | 28 | 28 | 28 | 35 | 35 | 35 |
|---|---|---|---|---|---|---|---|---|---|---|---|---|---|---|---|
| L | I | I | I | 4 | 4 | 4 | 4 | 4 | 4 | 4 | 4 | 4 | 4 | 4 | 4 |
| **Genera** | **I.1** | **I.2** | **I.3** | **4.1** | **4.2** | **4.3** | **4.1** | **4.2** | **4.3** | **4.1** | **4.2** | **4.3** | **4.1** | **4.2** | **4.3** |
| OLB8 | 1.8 | 2.3 | 1.8 | 22 | 27 | 26 | 15 | 37 | 33 | 10 | 38 | 27 | 17 | 19 | 20 |
| AKYH767(NA) | 3.5 | 4.2 | 3.2 | 7 | 8 | 8 | 9 | 5 | 4 | 9.6 | 4.8 | 4.8 | 8.3 | 7.1 | 8.0 |
| SBR1031(NA) | 0.1 | 0.8 | 0.7 | 2 | 4 | 3 | 9.6 | 7.9 | 4.6 | 6.9 | 6.9 | 5.7 | 5.6 | 8.1 | 7.8 |
| Glutamicibacter | 0.0 | 0.0 | 0.0 | 1.8 | 1.6 | 1.2 | 1.8 | 1.4 | 1.7 | 4.1 | 1.1 | 3.6 | 7.0 | 2.5 | 2.4 |
| Leucobacter | 0.0 | 0.0 | 0.0 | 11 | 4.9 | 5.6 | 4.4 | 1.8 | 3.9 | 3.9 | 1.6 | 4.2 | 5.7 | 2.0 | 4.0 |
| env.OPS_17(NA) | 2 | 2.6 | 2.0 | 0.9 | 1.5 | 1.2 | 0.5 | 1.0 | 0.3 | 0.6 | 0.3 | 1.1 | 0.8 | 7.3 | 2.7 |
| Chitinophagaceae(NA) | 2.0 | 2.3 | 1.8 | 0.0 | 0.0 | 0.1 | 0.2 | 0.7 | 0.2 | 1.0 | 1.0 | 0.9 | 3.5 | 3.0 | 3.3 |
| Saccharimonadales(NA) | 1.1 | 0.8 | 0.9 | 14 | 5.7 | 7.0 | 4.2 | 1.3 | 3.2 | 4.2 | 1.7 | 5.3 | 2.6 | 2.6 | 4.0 |
| Ca. Competibacter | 2.4 | 2.7 | 2.5 | 1.6 | 1.3 | 2.0 | 5.6 | 3.1 | 3.2 | 5.4 | 3.2 | 3.6 | 2.5 | 2.2 | 3.6 |
| Thauera | 0.9 | 0.8 | 0.8 | 1.2 | 1.4 | 3.7 | 0.2 | 0.6 | 0.3 | 1.1 | 1.0 | 0.8 | 4.0 | 3.4 | 0.6 |
| 45597(NA) | 0.1 | 0.5 | 0.0 | 0.4 | 0.3 | 0.1 | 1.6 | 0.6 | 0.7 | 2.7 | 1.4 | 1.2 | 3.6 | 2.4 | 2.0 |
| Microscillaceae(NA) | 5.5 | 5.1 | 4.4 | 1.2 | 1.4 | 1.6 | 2.6 | 2.1 | 1.8 | 4.0 | 1.9 | 3.0 | 2.5 | 2.0 | 3.4 |
| Nakamurella | 0.0 | 0.0 | 0.0 | 0.0 | 0.0 | 0.0 | 0.0 | 0.0 | 0.5 | 1.1 | 0.3 | 1.3 | 2.4 | 2.7 | 2.3 |
| Ferruginibacter | 2.0 | 1.8 | 1.7 | 1.6 | 2.1 | 2.2 | 3.2 | 2.5 | 1.5 | 2.7 | 2.3 | 2.3 | 2.1 | 2.5 | 2.6 |
| Niabella | 0.0 | 0.0 | 0.1 | 0.9 | 1.0 | 1.4 | 1.0 | 1.4 | 1.5 | 1.8 | 1.8 | 1.4 | 3.1 | 1.4 | 1.9 |
| Kouleothrix | 0.6 | 0.6 | 0.8 | 0.0 | 0.4 | 0.4 | 3.6 | 1.9 | 2.1 | 0.9 | 1.9 | 2.4 | 0.8 | 2.0 | 3.0 |
| OLB17 | 2 | 2.6 | 1.9 | 1 | 1.4 | 0.7 | 1.9 | 1.6 | 1.0 | 2.3 | 1.5 | 1.8 | 2.3 | 1.6 | 1.6 |
| Caldilineaceae(NA) | 0.5 | 0.4 | 0.5 | 2.1 | 1.1 | 1.4 | 4.2 | 1.8 | 1.7 | 2.7 | 2.0 | 2.2 | 1.2 | 1.6 | 1.9 |
| Gemmataceae(NA) | 1 | 0.0 | 0.8 | 0.2 | 0.0 | 0.3 | 1.1 | 0.5 | 0.8 | 1.7 | 1.2 | 1.2 | 1.2 | 0.9 | 1.9 |
| Plasticicumulans | 0.4 | 0.3 | 0.3 | 0.9 | 0.9 | 0.5 | 0.0 | 0.4 | 0.1 | 2.8 | 1.7 | 0.0 | 2.5 | 1.1 | 0.0 |

| d | 0 | 0 | 0 | 14 | 14 | 14 | 21 | 21 | 21 | 28 | 28 | 28 | 35 | 35 | 35 |
|---|---|---|---|---|---|---|---|---|---|---|---|---|---|---|---|
| L | I | I | I | 5 | 5 | 5 | 5 | 5 | 5 | 5 | 5 | 5 | 5 | 5 | 5 |
| **Genera** | **I.1** | **I.2** | **I.3** | **5.1** | **5.2** | **5.3** | **5.1** | **5.2** | **5.3** | **5.1** | **5.2** | **5.3** | **5.1** | **5.2** | **5.3** |
| OLB8 | 1.8 | 2.3 | 1.8 | 23 | 18 | 24 | 31 | 26 | 37 | 19 | 7 | 32 | 17 | 8 | 12 |
| SBR1031(NA) | 0.1 | 0.8 | 0.7 | 3 | 6 | 5 | 7 | 7 | 7 | 6.9 | 12 | 4.9 | 7.2 | 11 | 10 |
| AKYH767(NA) | 3.5 | 4.2 | 3.2 | 7 | 8 | 10 | 5.3 | 6.0 | 3.3 | 5.5 | 6.1 | 4.5 | 5.3 | 4.8 | 7.4 |
| Leucobacter | 0.0 | 0.0 | 0.0 | 10 | 8.9 | 3.8 | 4.5 | 3.2 | 2.1 | 5.4 | 4.1 | 2.6 | 5.5 | 3.2 | 4.4 |
| Thauera | 0.9 | 0.8 | 0.8 | 1 | 2.1 | 2.0 | 1.8 | 1.1 | 0.4 | 1.1 | 1.3 | 0.6 | 4.9 | 5.5 | 1.4 |
| Saccharimonadales(NA) | 1 | 0.8 | 0.9 | 12 | 12 | 6.5 | 4.3 | 6.2 | 4.3 | 6.2 | 5.0 | 3.1 | 6.5 | 2.3 | 2.9 |
| Ca. Competibacter | 2.4 | 2.7 | 2.5 | 1.7 | 1.1 | 2.0 | 2.7 | 3.3 | 2.3 | 4.2 | 4.8 | 2.3 | 3.2 | 3.8 | 4.1 |
| Kouleothrix | 0.6 | 0.6 | 0.8 | 0 | 0.7 | 1.7 | 0.8 | 3.2 | 2.6 | 2.6 | 3.6 | 2.8 | 2.2 | 3.8 | 3.3 |
| Microscillaceae(NA) | 5.5 | 5.1 | 4.4 | 1.5 | 1.6 | 1.3 | 2.5 | 2.2 | 1.5 | 2.9 | 3.0 | 2.1 | 2.9 | 3.0 | 3.3 |
| Glutamicibacter | 0.0 | 0.0 | 0.0 | 0.7 | 1.5 | 1.3 | 1.4 | 2.0 | 0.6 | 2.9 | 3.8 | 2.3 | 2.6 | 3.8 | 2.5 |
| 45597(NA) | 0.1 | 0.5 | 0.0 | 0.2 | 0.0 | 0.5 | 1.2 | 0.9 | 0.9 | 1.2 | 2.0 | 1.8 | 1.4 | 3.3 | 3.3 |
| Nakamurella | 0.0 | 0.0 | 0.0 | 0.0 | 0.0 | 0.0 | 0.0 | 0.5 | 0.3 | 1.4 | 2.1 | 1.0 | 2.1 | 2.9 | 2.7 |
| Saccharimonadaceae(NA) | 0.0 | 0.1 | 0.2 | 0.0 | 0.0 | 0.2 | 0.8 | 0.7 | 1.1 | 1.9 | 2.4 | 1.0 | 2.4 | 2.7 | 2.2 |
| Niabella | 0.0 | 0.0 | 0.1 | 1.7 | 2.0 | 1.2 | 2.1 | 1.7 | 1.1 | 2.6 | 1.2 | 1.4 | 2.6 | 1.8 | 2.4 |
| Ferruginibacter | 2.0 | 1.8 | 1.7 | 1.7 | 1.6 | 2.1 | 2.0 | 2.2 | 1.7 | 2.5 | 2.0 | 1.9 | 2.3 | 2.1 | 2.0 |
| Caldilineaceae(NA) | 0.5 | 0.4 | 0.5 | 1.7 | 2.0 | 1.9 | 2.4 | 2.2 | 1.7 | 2.3 | 2.5 | 2.0 | 1.5 | 2.1 | 2.0 |
| Chitinophagaceae(NA) | 2 | 2.3 | 1.8 | 0 | 0.1 | 0.0 | 0.2 | 0.3 | 0.3 | 0.5 | 0.7 | 0.8 | 0.7 | 1.4 | 3.2 |
| Plasticicumulans | 0.4 | 0.3 | 0.3 | 1.0 | 1.1 | 0.7 | 0.6 | 0.5 | 0.3 | 0.5 | 0.4 | 0.0 | 1.4 | 2.1 | 1.8 |
| Diaphorobacter | 0 | 0.0 | 0.0 | 5.0 | 2.5 | 2.4 | 0.5 | 1.4 | 1.5 | 2.5 | 2.0 | 3.7 | 2.1 | 1.8 | 1.0 |
| OLB17 | 2.1 | 2.6 | 1.9 | 1.0 | 1.5 | 1.8 | 1.8 | 1.7 | 1.5 | 1.8 | 1.7 | 1.8 | 1.3 | 1.5 | 2.1 |

| d | 0 | 0 | 0 | 14 | 14 | 14 | 21 | 21 | 21 | 28 | 28 | 28 | 35 | 35 | 35 |
|---|---|---|---|---|---|---|---|---|---|---|---|---|---|---|---|
| L | I | I | I | 6 | 6 | 6 | 6 | 6 | 6 | 6 | 6 | 6 | 6 | 6 | 6 |
| **Genera** | **I.1** | **I.2** | **I.3** | **6.1** | **6.2** | **6.3** | **6.1** | **6.2** | **6.3** | **6.1** | **6.2** | **6.3** | **6.1** | **6.2** | **6.3** |
| SBR1031(NA) | 0.1 | 0.8 | 0.7 | 3 | 4 | 3 | 7 | 6 | 4 | 10 | 10 | 9 | 14 | 12 | 14 |
| OLB8 | 1.8 | 2.3 | 1.8 | 18 | 21 | 29 | 32 | 32 | 23 | 24 | 21 | 23 | 14 | 14 | 10 |
| AKYH767(NA) | 3.5 | 4.2 | 3.2 | 5 | 5 | 7 | 2.7 | 6.2 | 7.9 | 4.6 | 4.2 | 5.2 | 6.9 | 5.4 | 6.0 |
| Saccharimonadales(NA) | 1.1 | 0.8 | 0.9 | 13 | 12 | 5.2 | 2.4 | 4.1 | 4.7 | 5.1 | 6.9 | 4.0 | 3.1 | 7.7 | 3.1 |
| Ca. Competibacter | 2.4 | 2.7 | 2.5 | 1 | 1.8 | 1.2 | 2.4 | 1.4 | 2.2 | 3.5 | 3.2 | 3.4 | 1.9 | 5.6 | 4.9 |
| Nakamurella | 0 | 0.0 | 0.0 | 0.0 | 0.0 | 0.0 | 0.5 | 0.7 | 0.3 | 1.1 | 1.3 | 1.3 | 3.8 | 2.9 | 3.5 |
| Kouleothrix | 0.6 | 0.6 | 0.8 | 1.6 | 0.5 | 0.5 | 2.0 | 0.6 | 0.9 | 2.4 | 2.5 | 2.7 | 2.7 | 3.3 | 3.8 |
| 45597(NA) | 0.1 | 0.5 | 0.0 | 0 | 0.2 | 0.2 | 1.1 | 0.6 | 0.2 | 1.8 | 1.6 | 1.5 | 3.5 | 2.2 | 3.6 |
| Glutamicibacter | 0.0 | 0.0 | 0.0 | 3.9 | 3.3 | 2.8 | 2.5 | 0.6 | 1.8 | 0.5 | 2.8 | 4.2 | 0.1 | 2.4 | 4.6 |
| Leucobacter | 0.0 | 0.0 | 0.0 | 9.8 | 11 | 4.1 | 2.4 | 4.3 | 4.4 | 1.6 | 3.5 | 1.5 | 1.6 | 3.7 | 1.7 |
| Niabella | 0.0 | 0.0 | 0.1 | 1.5 | 1.4 | 1.2 | 1.5 | 1.0 | 1.2 | 1.5 | 1.0 | 1.4 | 3.5 | 1.8 | 1.5 |
| Chitinophagaceae(NA) | 2.0 | 2.3 | 1.8 | 0.0 | 0.0 | 0.0 | 0.5 | 0.3 | 0.0 | 0.9 | 0.9 | 0.9 | 2.5 | 2.0 | 2.2 |
| Actinomycetaceae(NA) | 0.0 | 0.0 | 0.0 | 1.6 | 0.0 | 1.2 | 2.1 | 1.7 | 0.0 | 1.3 | 2.1 | 4.0 | 0.8 | 1.5 | 3.2 |
| OLB17 | 2.1 | 2.6 | 1.9 | 0.9 | 0.9 | 0.4 | 1.1 | 1.6 | 1.2 | 1.7 | 2.0 | 1.3 | 1.8 | 1.9 | 1.7 |
| Ferruginibacter | 2.0 | 1.8 | 1.7 | 1.4 | 1.9 | 1.3 | 1.7 | 1.8 | 2.0 | 1.8 | 2.0 | 1.9 | 1.7 | 2.1 | 1.5 |
| Thauera | 0.9 | 0.8 | 0.8 | 1.4 | 3.1 | 3.8 | 1.0 | 0.7 | 1.2 | 1.0 | 1.7 | 1.7 | 1.6 | 2.6 | 1.2 |
| Microscillaceae(NA) | 5 | 5.1 | 4.4 | 0 | 0.4 | 0.5 | 0.3 | 0.7 | 1.6 | 1.0 | 0.7 | 0.7 | 1.8 | 1.7 | 1.5 |
| Saccharimonadaceae(NA) | 0.0 | 0.1 | 0.2 | 0.9 | 0.9 | 0.2 | 0.3 | 0.9 | 0.2 | 1.5 | 2.9 | 1.5 | 1.0 | 1.9 | 1.6 |
| Rubrivivax | 0 | 0.0 | 0.0 | 0.3 | 0.0 | 0.0 | 0.6 | 0.8 | 0.0 | 0.7 | 0.8 | 0.0 | 1.7 | 1.4 | 1.3 |
| env.OPS_17(NA) | 2.2 | 2.6 | 2.0 | 0.3 | 0.5 | 0.1 | 1.0 | 0.5 | 0.8 | 2.4 | 2.0 | 1.7 | 0.7 | 2.5 | 1.1 |

| d | 0 | 0 | 0 | 14 | 14 | 14 | 21 | 21 | 21 | 28 | 28 | 28 | 35 | 35 | 35 |
|---|---|---|---|---|---|---|---|---|---|---|---|---|---|---|---|
| L | I | I | I | 7 | 7 | 7 | 7 | 7 | 7 | 7 | 7 | 7 | 7 | 7 | 7 |
| **Genera** | **I.1** | **I.2** | **I.3** | **7.1** | **7.2** | **7.3** | **7.1** | **7.2** | **7.3** | **7.1** | **7.2** | **7.3** | **7.1** | **7.2** | **7.3** |
| Glutamicibacter | 0.0 | 0.0 | 0.0 | 10 | 7 | 9 | 10 | 3 | 9 | 1 | 5 | 9 | 1 | 4 | 25 |
| SBR1031(NA) | 0.1 | 0.8 | 0.7 | 4 | 3 | 3 | 4 | 5 | 4 | 7 | 6 | 7 | 9 | 9 | 7 |
| Saccharimonadales(NA) | 1.1 | 0.8 | 0.9 | 17 | 12 | 11 | 9.9 | 8.1 | 13 | 4.9 | 5.8 | 7.0 | 7.8 | 9.5 | 5.3 |
| Thauera | 0.9 | 0.8 | 0.8 | 2 | 4 | 5.0 | 3.7 | 6.0 | 6.1 | 11 | 11 | 6.9 | 9.3 | 5.8 | 5.9 |
| Sediminibacterium | 0.0 | 0.0 | 0.0 | 0 | 0.0 | 0.0 | 0.0 | 0.0 | 0.0 | 0.9 | 0.9 | 0.4 | 4.9 | 4.0 | 3.7 |
| Actinomycetaceae(NA) | 0 | 0.0 | 0.0 | 1.6 | 1.4 | 1.4 | 3.6 | 4.2 | 2.6 | 2.5 | 3.8 | 3.6 | 2.7 | 3.7 | 3.9 |
| OLB8 | 1.8 | 2.3 | 1.8 | 9.3 | 8.2 | 9.5 | 13 | 14 | 10 | 15 | 13 | 13 | 3.1 | 2.7 | 4.5 |
| Saccharimonadaceae(NA) | 0.0 | 0.1 | 0.2 | 2 | 1.0 | 0.9 | 3.5 | 2.1 | 2.5 | 2.3 | 3.3 | 2.3 | 2.8 | 4.8 | 1.9 |
| Diaphorobacter | 0.0 | 0.0 | 0.0 | 1.3 | 1.5 | 1.9 | 4.1 | 2.6 | 2.6 | 4.8 | 3.5 | 5.1 | 3.0 | 2.7 | 3.0 |
| Leucobacter | 0.0 | 0.0 | 0.0 | 14 | 20 | 13 | 1.8 | 2.7 | 8.6 | 2.0 | 1.4 | 2.7 | 3.4 | 2.4 | 2.4 |
| Alicycliphilus | 0.0 | 0.0 | 0.0 | 0.5 | 0.9 | 0.9 | 1.3 | 1.4 | 2.0 | 1.7 | 2.4 | 1.3 | 2.4 | 3.2 | 2.5 |
| Gemmatimonas | 0.0 | 0.0 | 0.0 | 0.0 | 0.2 | 0.0 | 0.7 | 0.0 | 0.3 | 1.0 | 0.9 | 1.0 | 2.6 | 2.8 | 2.5 |
| Pseudochrobactrum | 0.0 | 0.0 | 0.0 | 0.0 | 0.0 | 0.0 | 1.3 | 1.6 | 1.1 | 2.2 | 1.4 | 1.5 | 2.6 | 1.9 | 2.6 |
| Niabella | 0.0 | 0.0 | 0.1 | 1.8 | 2.7 | 1.2 | 2.4 | 2.0 | 1.6 | 2.1 | 2.1 | 1.8 | 2.1 | 2.3 | 2.0 |
| Burkholderiaceae(NA) | 1.0 | 1.1 | 1.0 | 0.0 | 0.1 | 0.1 | 0.0 | 0.0 | 0.0 | 0.4 | 0.4 | 0.3 | 2.7 | 2.0 | 1.2 |
| AKYG1722(NA) | 0.0 | 0.0 | 0.0 | 0.0 | 0.0 | 0.0 | 0.6 | 0.4 | 0.0 | 0.9 | 0.0 | 0.5 | 2.2 | 1.8 | 1.4 |
| Ca. Competibacter | 2 | 2.7 | 2.5 | 0 | 0.6 | 0.6 | 0.0 | 0.4 | 0.4 | 1.0 | 1.2 | 1.5 | 1.9 | 1.5 | 1.9 |
| Bosea | 0.0 | 0.0 | 0.0 | 1.1 | 0.6 | 1.1 | 0.8 | 2.0 | 2.5 | 1.7 | 1.1 | 1.9 | 2.4 | 1.3 | 0.8 |
| Pseudaminobacter | 0 | 0.0 | 0.0 | 0.0 | 0.0 | 0.0 | 0.0 | 0.1 | 0.2 | 0.9 | 0.8 | 1.1 | 1.7 | 1.3 | 1.5 |
| Fluviicola | 0.0 | 0.0 | 0.0 | 0.0 | 0.1 | 0.2 | 1.8 | 1.0 | 0.5 | 0.7 | 2.0 | 0.3 | 0.1 | 4.4 | 0.0 |

**Fig. S4.** Relative abundance heat maps for the top 20 genera, obtained through 16 rRNA amplicon sequencing. Each panel corresponds to a level of disturbance and columns correspond to a replicate reactor across all time points sampled. Changes in bacterial abundances with time and disturbance levels display differential succession. NA: not assigned at genus level (last assigned taxonomic category shown).

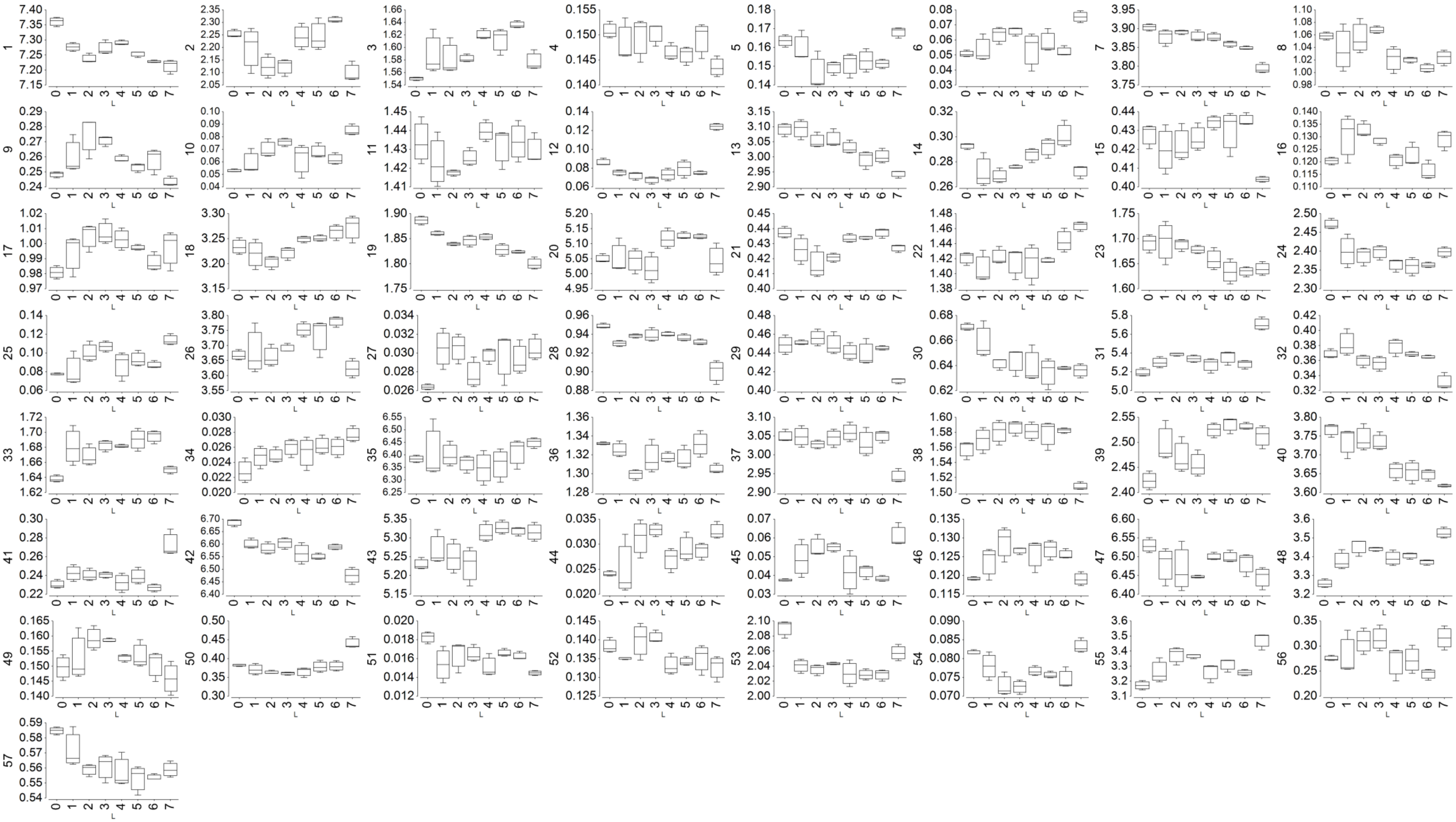


**Fig. S5.** Box plots of relative abundances (%) for genotypic trait complex categories of the IP2G database (gene ontologies) across reactors at different levels of disturbance (n = 3). Titles of y-axes represent each genotypic category, see references and univariate test results in Table S2.

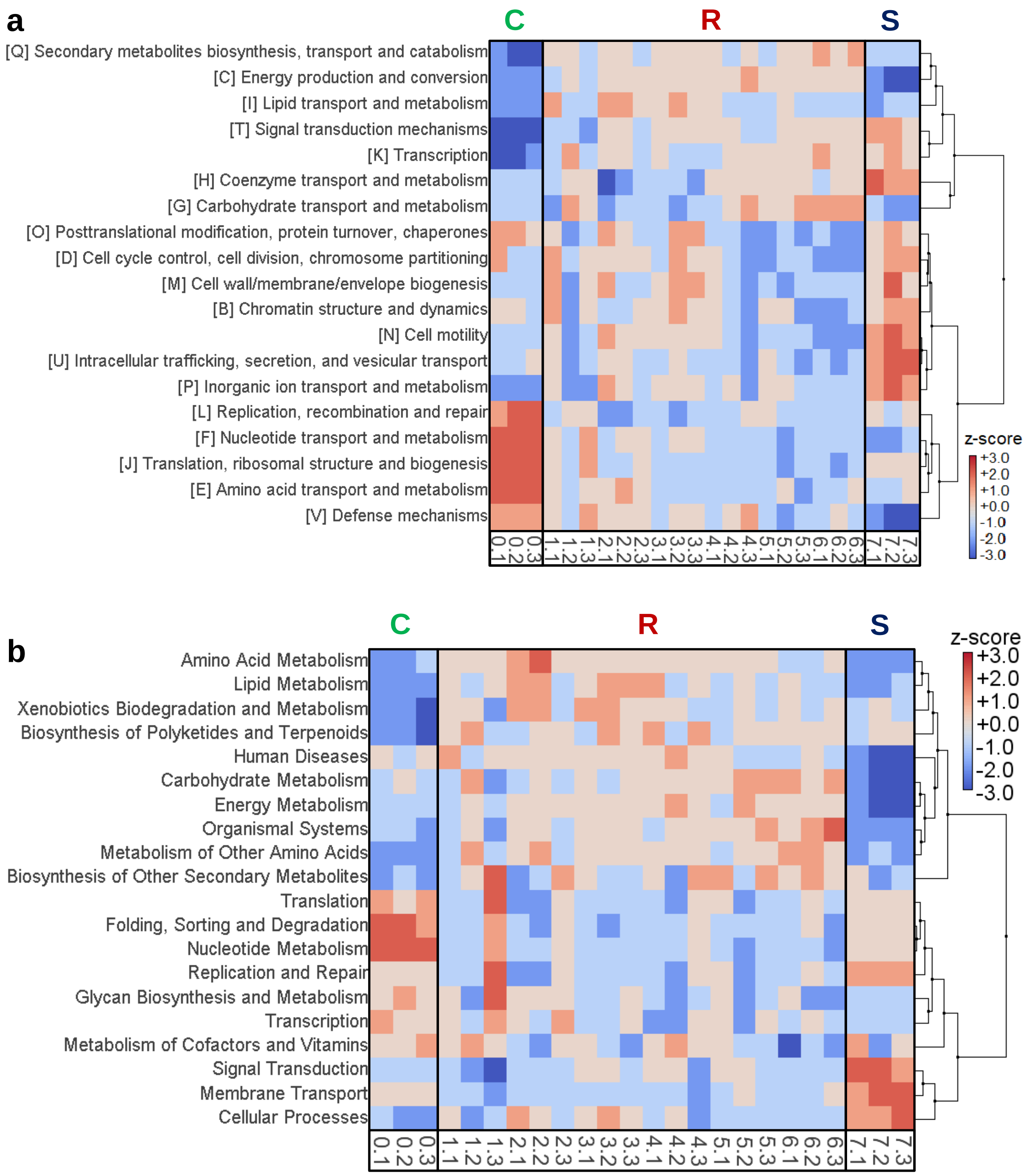
a
C
R
S
[Q] Secondary metabolites biosynthesis, transport and catabolism
[C] Energy production and conversion
[I] Lipid transport and metabolism
[T] Signal transduction mechanisms
[K] Transcription
[H] Coenzyme transport and metabolism
[G] Carbohydrate transport and metabolism
[O] Posttranslational modification, protein turnover, chaperones
[D] Cell cycle control, cell division, chromosome partitioning
[M] Cell wall/membrane/envelope biogenesis
[B] Chromatin structure and dynamics
[N] Cell motility
[U] Intracellular trafficking, secretion, and vesicular transport
[P] Inorganic ion transport and metabolism
[L] Replication, recombination and repair
[F] Nucleotide transport and metabolism
[J] Translation, ribosomal structure and biogenesis
[E] Amino acid transport and metabolism
[V] Defense mechanisms
0.1 0.2 0.3 1.1 1.2 1.3 2.1 2.2 2.3 3.1 3.2 3.3 4.1 4.2 4.3 5.1 5.2 5.3 6.1 6.2 6.3 7.1 7.2 7.3
z-score
+3.0
+2.0
+1.0
+0.0
-1.0
-2.0
-3.0
b
C
R
S
Amino Acid Metabolism
Lipid Metabolism
Xenobiotics Biodegradation and Metabolism
Biosynthesis of Polyketides and Terpenoids
Human Diseases
Carbohydrate Metabolism
Energy Metabolism
Organismal Systems
Metabolism of Other Amino Acids
Biosynthesis of Other Secondary Metabolites
Translation
Folding, Sorting and Degradation
Nucleotide Metabolism
Replication and Repair
Glycan Biosynthesis and Metabolism
Transcription
Metabolism of Cofactors and Vitamins
Signal Transduction
Membrane Transport
Cellular Processes
0.1 0.2 0.3 1.1 1.2 1.3 2.1 2.2 2.3 3.1 3.2 3.3 4.1 4.2 4.3 5.1 5.2 5.3 6.1 6.2 6.3 7.1 7.2 7.3
z-score
+3.0
+2.0
+1.0
+0.0
-1.0
-2.0
-3.0

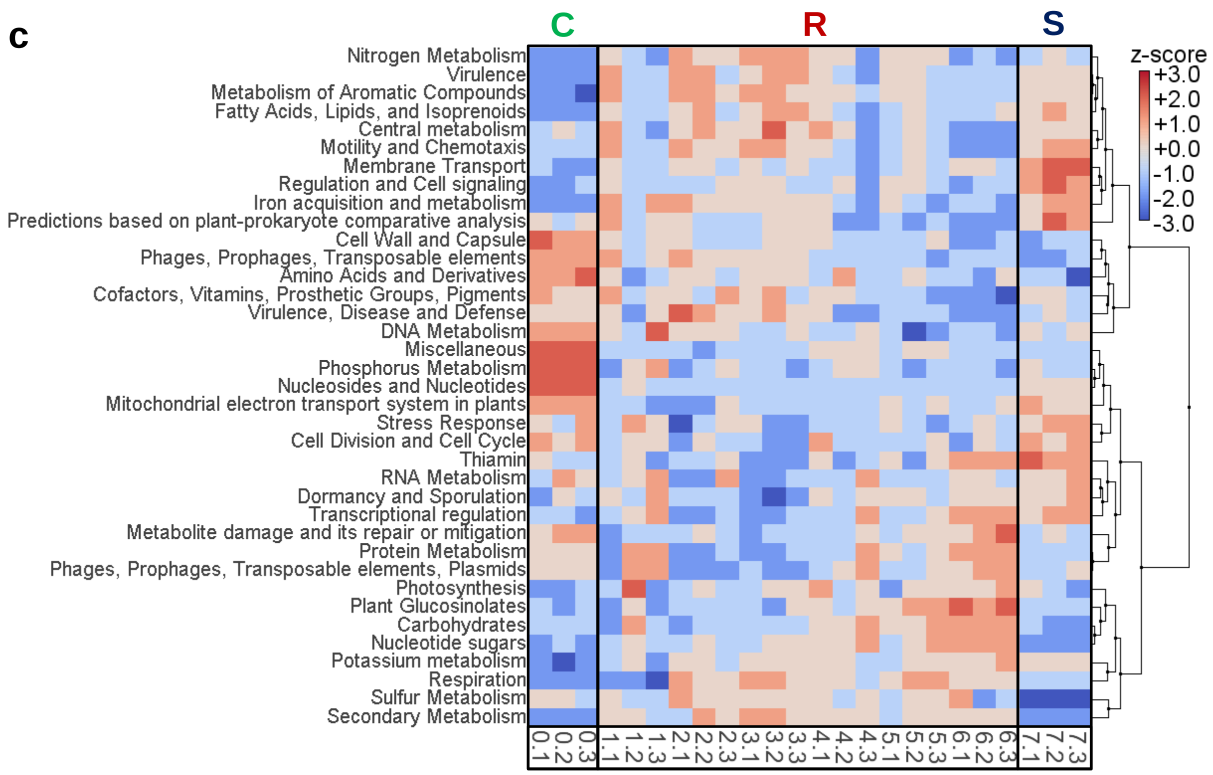


**Fig. S6.** Heat map of genotypic potential traits after mapping metagenomes with (**a**) COG, (**b**) KEGG, and (**c**) SEED databases using MEGAN. Functional capacity was classified using different trait complexes, including only those with more than 10,000 reads assigned across all reactors. Clustering was applied to differentiate groups of trait complexes and disturbance levels. Column legend represents disturbance level and replicate reactor number (n = 24). Rectangles highlight taxa groups prevailing at different CSR life-history strategies at the community-level: C, competitors (L0); R, ruderals (L1-6); S, stress-tolerants (L7).

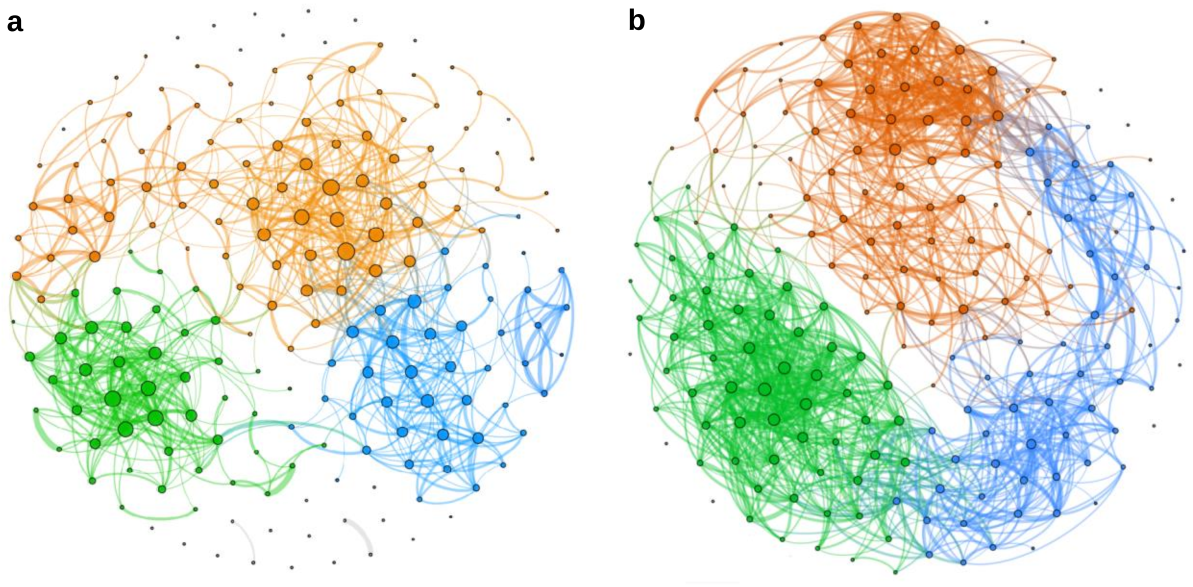

**Fig. S7.** Correlation networks display separated clusters based on node modularity for bacterial taxa and genes. (**a**) Top 200 amplicon sequence variants (ASVs). (**b**) Top 200 individual genes from IP2G database. Clusters are colored by modularity class, with green nodes prevailing in undisturbed (L0) reactors, orange nodes in intermediately disturbed reactors (L1-6), and blue nodes in press-disturbed reactors (L7). Only significantly strong Pearson's correlations ($r_a \geq 0.50$, $r_b \geq 0.60$) were employed. For each panel, edge thickness represents correlation strength and node size represents degree.

### Supporting Tables

**Table S1.** Community-level traits of process performance and productivity on day 35

| Community Trait[†] | Disturbance Levels* | | | | | | | | $P_{BH}$[§] |
|---|---|---|---|---|---|---|---|---|---|
| | 0 | 1 | 2 | 3 | 4 | 5 | 6 | 7 | |
| %COD rem[¶] | 99.5 (0.5) | 84.9 (0.7) | 86.2 (3.6) | 84.7 (0.7) | 96.1 (3.9) | 94.5 (2.3) | 98.4 (0.5) | 97.4 (0.7) | $1.20 \times 10^{-6}$ |
| %3-CA rem[¶] | - | 100 (0) | 100 (0) | 100 (0) | 100 (0) | 100 (0) | 100 (0) | 100 (0) | - |
| %$PO_4$-P rem[¶] | 13.4 (1.4) | 12.0 (1.1) | 12.6 (0.7) | 13.4 (0.5) | 11.6 (5.8) | 11.7 (0.9) | 17.3 (3.3) | 16.2 (1.6) | 0.1495 |
| %$NH_3$-N rem[¶#] | 49.5 (5.1) | 28.9 (11.4) | 10.7 (27.4) | 31.5 (2.6) | -46.8 (25.6) | -37.4 (18.5) | -70.5 (11.1) | -81.4 (7.8) | $1.60 \times 10^{-6}$ |
| $NO_2$-N [mg/L] | 0 (0) | 72.4 (3.7) | 59.2 (18.8) | 69.7 (3.1) | 15.8 (20.7) | 24.6 (18.3) | 0.07 (0.12) | 0 (0) | $1.54 \times 10^{-6}$ |
| $NO_3$-N [mg/L] | 61.4 (6.6) | 3.3 (3.9) | 0.4 (0.3) | 0.3 (0.4) | 0.2 (0.1) | 0.3 (0.2) | 0.03 (0.04) | 0 (0) | $3.21 \times 10^{-13}$ |
| TSS [g/L] | 6.8 (0.4) | 7.5 (0.1) | 6.5 (0.3) | 6.5 (0.2) | 6.6 (0.4) | 6.3 (0.7) | 6.2 (0.2) | 5.2 (0.2) | 0.0002 |
| VSS [g/L] | 5.9 (0.3) | 6.6 (0.2) | 5.8 (0.3) | 5.8 (0.2) | 5.9 (0.4) | 5.7 (0.7) | 5.5 (0.2) | 4.7 (0.0) | 0.0002 |
| %VSS:TSS | 86.1 (1.4) | 87.8 (1.4) | 89.4 (1.0) | 89.3 (1.7) | 88.9 (1.1) | 89.7 (1.3) | 89.6 (0.1) | 90.6 (2.0) | 0.3092 |
| %GC content | 61.0 (0.0) | 60.7 (0.9) | 60.0 (0.4) | 60.5 (0.4) | 61.3 (0.2) | 61.5 (0.4) | 61.8 (0.2) | 62.3 (0.2) | 0.0020 |

[*] Average values (n = 3), including standard deviation of the mean in parentheses.

[†] Ecosystem function or community-level trait

[‡] Welch's ANOVA test P-value

[§] Benjamini-Hochberg corrected P-value at FDR = 10%

[¶] Removal of the indicated compound (percentage based on the feed input)

[||] ANOVA test P-value, as at least one group had zero variance

[#] Negative values represent ammonia accumulation due to nitrification inhibition

**Table S2.** Welch's ANOVA tests comparing each of the trait complexes from IP2G database (>10000 reads), across different disturbance levels adjusted at a FDR of 10%. CSR categories were assigned as described in the main text.

| # | Gene Ontologies Number | Trait Complex Classification (IP2G) | Welch P-value | Benjamini-Hochberg P-value | Benjamini-Hochberg significance | CSR category |
|---|---|---|---|---|---|---|
| 1 | GO:0009058 | biosynthetic process | 0.00020 | 0.00158 | significant | C |
| 2 | GO:0005975 | carbohydrate metabolic process | 0.00123 | 0.00334 | significant | R |
| 3 | GO:0009056 | catabolic process | 0.00002 | 0.00104 | significant | R |
| 4 | GO:0007154 | cell communication | 0.10141 | 0.10510 | not significant | CSR |
| 5 | GO:0051301 | cell division | 0.00705 | 0.01057 | significant | SC |
| 6 | GO:0048870 | cell motility | 0.00293 | 0.00521 | significant | S |
| 7 | GO:0006520 | cellular amino acid metabolic process | 0.00051 | 0.00211 | significant | C |
| 8 | GO:0016043 | cellular component organization | 0.00057 | 0.00215 | significant | CR |
| 9 | GO:0045333 | cellular respiration | 0.00192 | 0.00406 | significant | R |
| 10 | GO:0006935 | chemotaxis | 0.00043 | 0.00206 | significant | SR |
| 11 | GO:0051186 | cofactor metabolic process | 0.02683 | 0.03186 | significant | CR |
| 12 | GO:0000746 | conjugation | 0.00004 | 0.00104 | significant | S |
| 13 | GO:0006259 | DNA metabolic process | 0.00139 | 0.00345 | significant | CR |
| 14 | GO:0022900 | electron transport chain | 0.00262 | 0.00498 | significant | R |
| 15 | GO:0006091 | generation of precursor metabolites and energy | 0.00014 | 0.00158 | significant | R |
| 16 | GO:0042592 | homeostatic process | 0.00931 | 0.01277 | significant | SR |
| 17 | GO:0006629 | lipid metabolic process | 0.03571 | 0.04155 | significant | R |
| 18 | GO:0006807 | nitrogen compound metabolic process | 0.03686 | 0.04202 | significant | SR |
| 19 | GO:0009117 | nucleotide metabolic process | 0.00024 | 0.00158 | significant | C |
| 20 | GO:0055114 | oxidation-reduction process | 0.00876 | 0.01277 | significant | R |
| 21 | GO:0016310 | phosphorylation | 0.01073 | 0.01390 | significant | CR |
| 22 | GO:0019538 | protein metabolic process | 0.00318 | 0.00533 | significant | SR |
| 23 | GO:0006950 | response to stress | 0.01601 | 0.01942 | significant | CR |
| 24 | GO:0016070 | RNA metabolic process | 0.00174 | 0.00381 | significant | C |
| 25 | GO:0007165 | signal transduction | 0.00119 | 0.00334 | significant | SR |
| 26 | GO:0044281 | small molecule metabolic process | 0.00490 | 0.00755 | significant | R |
| 27 | GO:0043934 | sporulation | 0.00941 | 0.01277 | significant | SR |
| 28 | GO:0006790 | sulfur compound metabolic process | 0.00478 | 0.00755 | significant | C |
| 29 | GO:0006351 | transcription, DNA-templated | 0.00011 | 0.00157 | significant | CR |
| 30 | GO:0006412 | translation | 0.00022 | 0.00158 | significant | CR |
| 31 | GO:0006810 | transport | 0.00244 | 0.00489 | significant | S |
| 32 | GO:0032196 | transposition | 0.05720 | 0.06038 | significant | R |
| 33 | GO:0048037 | cofactor binding | 0.00029 | 0.00158 | significant | R |
| 34 | GO:0030234 | enzyme regulator activity | 0.17838 | 0.17838 | not significant | CSR |
| 35 | GO:0016787 | hydrolase activity | 0.16529 | 0.16824 | not significant | CSR |
| 36 | GO:0016853 | isomerase activity | 0.00171 | 0.00381 | significant | CR |
| 37 | GO:0016874 | ligase activity | 0.01007 | 0.01335 | significant | R |
| 38 | GO:0016829 | lyase activity | 0.00008 | 0.00152 | significant | R |
| 39 | GO:0046872 | metal ion binding | 0.00417 | 0.00679 | significant | R |
| 40 | GO:0003676 | nucleic acid binding | 0.00030 | 0.00158 | significant | CR |
| 41 | GO:0001071 | nucleic acid binding transcription factor activity | 0.03851 | 0.04304 | significant | S |
| 42 | GO:0000166 | nucleotide binding | 0.00083 | 0.00264 | significant | C |
| 43 | GO:0016491 | oxidoreductase activity | 0.00909 | 0.01277 | significant | SR |
| 44 | GO:0000156 | phosphorelay response regulator activity | 0.00129 | 0.00335 | significant | SR |
| 45 | GO:0004872 | receptor activity | 0.00082 | 0.00264 | significant | SR |
| 46 | GO:0000988 | transcription factor activity, protein binding | 0.00074 | 0.00263 | significant | R |
| 47 | GO:0016740 | transferase activity | 0.00314 | 0.00533 | significant | CR |
| 48 | GO:0005215 | transporter activity | 0.00052 | 0.00211 | significant | SR |
| 49 | GO:0051082 | unfolded protein binding | 0.01377 | 0.01706 | significant | R |
| 50 | GO:0043190 | ATP-binding cassette (ABC) transporter complex | 0.00289 | 0.00521 | significant | S |
| 51 | GO:0005618 | cell wall | 0.00249 | 0.00489 | significant | C |
| 52 | GO:0005694 | chromosome | 0.05022 | 0.05504 | significant | CR |
| 53 | GO:0005737 | cytoplasm | 0.01264 | 0.01601 | significant | SC |
| 54 | GO:0042575 | DNA polymerase complex | 0.00121 | 0.00334 | significant | SC |
| 55 | GO:0016020 | membrane | 0.00162 | 0.00381 | significant | SR |
| 56 | GO:0042597 | periplasmic space | 0.05332 | 0.05735 | significant | SR |
| 57 | GO:0005840 | ribosome | 0.00029 | 0.00158 | significant | CR |
